# Circular RNAs arising from synaptic host genes are modulated by SFPQ RNA-binding protein and increased during human neuronal differentiation

**DOI:** 10.1101/2022.10.05.510949

**Authors:** Michelle Watts, Marika Oksanen, Sanna Lejerkrans, Francesca Mastropasqua, Myriam Gorospe, Kristiina Tammimies

## Abstract

**Background:** Circular RNAs (circRNAs) are emerging as a key component of the complex neural transcriptome implicated in brain development. However, the specific expression patterns and functions of circRNAs in human neuronal differentiation have not been explored.

**Results:** Using total RNA sequencing analysis, we identified expressed circRNAs during the differentiation of human neuroepithelial stem (NES) cells into developing neurons and discovered that many circRNAs originated from host genes associated with synaptic function. Interestingly, when assessing population data, exons giving rise to circRNAs in our dataset had a higher frequency of genetic variants. Additionally, screening for RNA-binding protein sites identified enrichment of Splicing Factor Proline and Glutamine Rich (SFPQ) motifs in increased circRNAs, several of which were reduced by SFPQ knockdown and enriched in SFPQ ribonucleoprotein complexes.

**Conclusions:** Our study provides an in-depth characterisation of circRNAs in a human neuronal differentiation model and highlights SFPQ as both a regulator and binding partner of circRNAs elevated during neuronal maturation.

## Background

CircRNAs represent a unique class of RNA molecules formed following non-canonical back-splicing of exon-exon junctions (Cocquerelle *et al*, 1992; Nigro *et al*, 1991). These covalently closed, circularised RNAs lack 5’cap and 3’poly-A structures resulting in highly stable molecules, resistant to enzymatic degradation (Jeck *et al*, 2013; Memczak *et al*, 2013). Since their initial discovery, the circRNA transcriptome has expanded rapidly with the advent of sequencing approaches and bioinformatic tools designed to detect back-spliced junctions in RNA-sequencing (RNAseq) data (Memczak *et al*., 2013; Salzman *et al*, 2013; Salzman *et al*, 2012). Although once thought to be non-functional by-products of spurious splicing events, numerous functional roles of circRNAs have now been identified, including microRNA (miRNA) sponging (Memczak *et al*., 2013), transcriptional regulation (Conn *et al*, 2017; Pandey *et al*, 2020), RNA-binding protein function (Panda *et al*, 2017; Tsitsipatis *et al*, 2021), and modulation of protein interactions (Barbagallo *et al*, 2019; Du *et al*, 2016). These roles highlight the vast diversity of possible functions for circRNAs, most of which are unknown.

As is often observed for mechanisms such as microRNAs or splicing factors that diversify the mammalian transcriptome, circRNA species are highly enriched in the nervous system (Memczak *et al*., 2013; Rybak-Wolf *et al*, 2015; You *et al*, 2015). Spatio-temporal regulation of circRNA expression within the brain is also dynamic, differing across brain regions and developmental stages (Chen *et al*, 2019; Veno *et al*, 2015; You *et al*., 2015). Brain and neuronal expressed circRNAs have also been found to arise from back-splicing of linear host genes associated with synaptic functions and display specific enrichment within synaptosomes as well as regulation in response to neuronal activation (Gokool *et al*, 2020; You *et al*., 2015). Under various circumstances, this regulation appears independent of corresponding host genes and, in some cases, circRNA levels far exceed linear host RNA levels (Rybak-Wolf *et al*., 2015; You *et al*., 2015), underscoring the importance of precise modulation of circRNAs during neuronal development.

Accumulating evidence suggests that circRNAs are not only important for neuronal function but also for brain development. Differential expression of circRNAs has been found to occur during neuronal differentiation, both in mice and in cell culture models such as the human neuroblastoma cell line SH-SY5Y, with several specific circRNAs were found to have a regulatory role in this process (Hollensen *et al*, 2020; Li *et al*, 2022b; Suenkel *et al*, 2020; Yang *et al*, 2019; Yang *et al*, 2018). Additionally, a number of circRNAs have been identified as being dysregulated in Autism Spectrum Disorder (ASD) (Chen *et al*, 2020; Gokool *et al*., 2020), while a depletion of circRNAs has been observed in individuals with Schizophrenia (SCZ) (Mahmoudi *et al*, 2019). Currently, however, our understanding of circRNA regulation and function in human neurodevelopment remains largely unexplored.

While evaluation of circRNAs in the human brain is limited to post-mortem analyses, derivation of mature neurons from human induced pluripotent stem cells (hiPSCs) allows for modelling of neuronal differentiation in a human genetic background (Dolmetsch & Geschwind, 2011). Computational methods to detect back-spliced junctions in total RNAseq data also allow for the discovery of circRNAs without the need for specialised treatment of RNA samples as well as comparison with linear RNA expression from the same data (Hansen, 2018). We combined these methodologies to examine the circRNA landscape of hiPSC-derived neuroepithelial stem (NES) cells and developing neurons, revealing an increase in circRNAs during differentiation, arising from host genes with synaptic functions. CircRNA-forming exons were also found to harbour genetic variants at a higher rate in comparison to exons from the same host gene that were not circularized. In addition, we investigated the potential functionality of elevated circRNAs and identified an enrichment of binding motifs for Splicing Factor Proline and Glutamine Rich (SFPQ). Predicted circRNA targets of SFPQ were found to be depleted during neuronal differentiation following SFPQ knockdown including a novel circRNA from the Regulating Synaptic Membrane Exocytosis 1 (RIMS1) host gene that was specifically enriched by SFPQ immunoprecipitation above its corresponding linear isoform.

## Results

### Detection of robust circRNA production during NES differentiation from total RNAseq data

To analyse circRNAs in human neuronal differentiation, hiPSC-derived NES cells were differentiated as previously described (Becker *et al*, 2020). Total RNA was collected and sequenced from cells in their proliferative, neural progenitor state (NES) and two time points of differentiation. First, at day five (D5), when a neural identity has been adopted and neurite outgrowth/axon specification is underway, and second at day 28 (D28), when extensive neural networks, but not yet mature active connections, have been established **[Fig 1a]**. To reduce spurious circRNA discovery, three detection programs (CIRCexplorer2, CIRI, MapSplice) with reported low false positive rates were utilised (Hansen, 2018). A total of 60,531 detected circRNAs were filtered for minimum expression, leaving 6,540 circRNAs (∼10.8%) for downstream analysis. These were further filtered to include only circRNAs detected by at least two programs, which represented the majority of the expression filtered dataset (6,385; ∼97.6%) **[Fig 1b]**. Among these, 66% were known circRNAs reported in circBase (Glazar *et al*, 2014), 34% were novel circRNAs (not yet reported in circBase), and 8.5% of the dataset were ‘*de novo’* circRNAs that did not annotate to known splice sites. The entire filtered dataset of 6,385 circRNAs was derived from 2,743 unique host genes, with 1,377 genes producing two or more circRNA isoforms and 50 ‘hotspot’ genes that generated over ten circRNAs **[Additional File 2: Table S1a]**. All circRNAs analysed and reported in Additional File 2: Table S1 have been assigned an ID based on host gene name and back-spliced junction genomic locations. Where available, corresponding circBase IDs are provided. For simplicity, specific circRNAs referenced within this text are referred to only by host gene name with alphabetical indexing for referencing circRNA isoforms.

**Figure 1.**
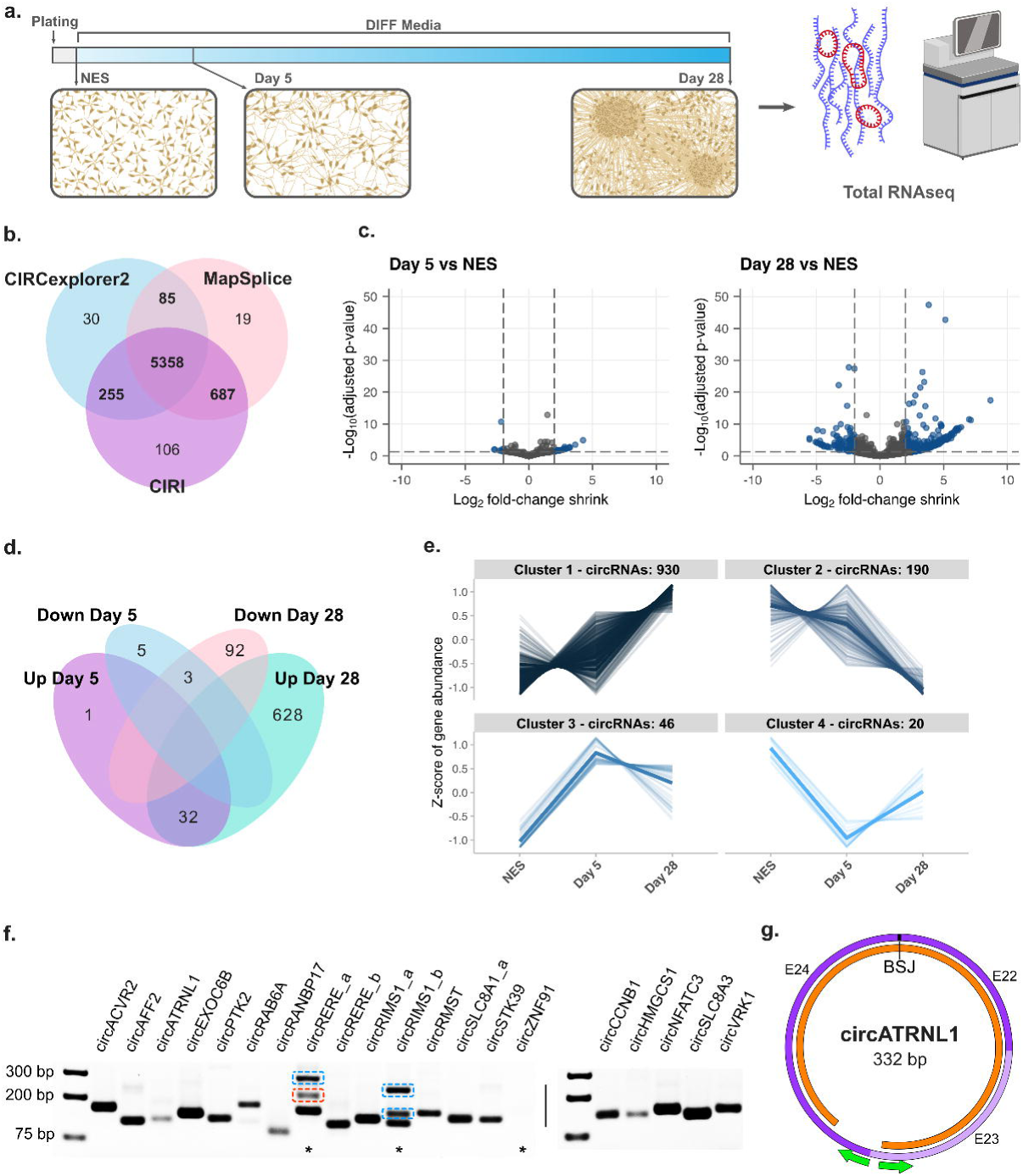
Detection and validation of circRNAs in human neuronal differentiation**. a.** Schematic representation of sample and data collection. **b.** Venn diagram of circRNAs detected from three different circRNA detection programs, numbers highlighted in bold indicate circRNAs included in downstream analysis. **c.** Volcano plots of differentially expressed circRNAs at D5 and D28 of differentiation compared to NES. **d.** Venn diagram of significantly dysregulated circRNAs (padj < 0.05 and LFC ≥|2|). **e.** Cluster analysis of circRNAs differentially expressed throughout differentiation. **f.** Detection of top differentially expressed circRNAs by PCR with divergent junction primers from RNAse R-treated samples. Samples from D28 were used to detect 15 increased circRNAs (left) and samples from NES cells were used for detection of 5 decreased circRNAs (right). Asterisks indicate reactions with additional bands or no product, blue highlighting indicates the product is an isoform of the intended circRNA target, red highlighting indicates PCR artifact. **g.** Example schematic of sanger sequencing validation of circRNA exon retention in *circATRNL1*. Exons are shown in purple, green arrows indicate primer location, and sequencing alignment is shown in orange.

Differentially expressed circRNAs (DEcircs) in NES differentiation were detected using DESeq2 with log fold-change (LFC) shrinkage estimation for increased stringency. We observed a predominant increase of circRNA expression with pairwise comparisons finding 33 circRNAs up and 8 circRNAs down at D5 compared to NES and 660 circRNAs up and 95 down at D28 compared to NES (p-adjusted <0.05, LFC ≥|2|) **[Fig 1c; Additional File 2: Table S1b-d]**. Almost all circRNAs (32/33) whose expression was increased at the early differentiation time point (D5) remained significantly increased at D28 **[Fig 1d]**. Clustering results using the likelihood ratio test, which incorporates all time points, similarly highlighted a primary cluster of circRNAs with a pattern of increasing abundance throughout differentiation **[Fig 1e; Additional File 2: Table S1e,f]**.

To validate whether detected circRNAs occurred endogenously and are differentially expressed during differentiation, we selected 20 DEcircs and designed divergent primer pairs to selectively amplify back-spliced junctions **[Additional File 1: Fig S1a]**. PCR amplification and gel electrophoresis found a clear, predominant band of expected size for 19/20 primer pairs in samples that were depleted of linear RNA by RNAse R treatment **[Fig 1f]**. No amplification was observed from primers designed for *de novo* circRNA *circZNF1* and primers for *circRERE_a_*and *circRIMS1_b* produced bands in addition to the expected product **[Fig 1f]**. Sanger sequencing confirmed that for *circRERE_a* primers, these corresponded to either PCR artifact or to the *circRERE_b* isoform, while primers for *circRIMS1_b* produced two additional *circRIMS1* isoforms that contained either one or two additional exons not annotated in NCBI **[Additional File 1: Fig S1b,c]**. These unannotated exons (UEs) mapped to the RIMS1 intronic region between exons 26 and 27 (UE-1 = 28-b; GRCh37-chr6:73005640-73005667, UE-2 = 90-b; GRCh37-chr6:73008851-73008880). Using quantitative PCR (qPCR) analysis of replicate samples, similar differential expression of circRNAs was also observed at D28 when compared to NES, particularly for increased circRNAs **[Additional File 1: Fig S1d]** and qPCR LFC values strongly correlated with RNAseq LFC values (R = 0.8, p = 1.2e-4). Finally, secondary divergent primer pairs were designed to confirm circRNA annotation and determine internal exon structure in circRNAs with non-consecutive back-spliced exons **[Additional File 1: Fig S2a]**. Among primer pairs which amplified enough product, all except *circRIMS1_a*, produced a single band. Sanger sequencing confirmed back-spliced exon identity in all circRNAs as well as internal exon retention, where relevant **[Fig 1g and Additional File 1: Fig S2b-m]**. A second larger product of the primer pair for *circRIMS1_a* (*circRIMS1_a2*) was found to also contain the 90-b UE-2 described above. To determine if these UEs are expressed in linear RIMS1 RNA, we designed primers flanking this region and performed PCR amplification on D28 cDNA. A single product was detected that we confirmed by Sanger sequencing excluded both UEs, indicating these exons may be circRNA specific **[Additional File 1: Fig S2n]**. Finally, as an additional validation, the dataset was compared with circRNAs detected by Gokool *et al*, which represents the largest dataset of human brain circRNAs (197 post-mortem samples), and ∼80% of circRNAs in our study were found to be represented. Taken together, these results indicate that our pipeline for circRNA detection from total RNAseq data was successful in identifying genuine circRNAs that are differentially expressed during neuronal differentiation.

### CircRNAs increased during NES differentiation arise from synaptic host genes

To gain insight into the host genes from which circRNAs are produced during differentiation, functional annotations associated with circRNA host genes were examined. As NES differentiation primarily increased circRNA expression, we initially tested host gene ontology enrichment among Cluster 1 circRNAs as well as positive gene set enrichment analysis (GSEA) of D28 vs NES DEcircs. This highlighted significantly enriched terms predominantly related to synaptic functions including synapse assembly, synaptic vesicle cycle and ion transport in addition to other neuronal development processes such as neuronal projection and cAMP signalling **[Fig 2a,b and Additional File 3: Table S2a,b]**. While similar pathways were identified as positively enriched from GSEA of D5 vs NES, only one reached significance (GO:0051962-Positive regulation of nervous system development; p-adjusted <0.05). Of interest, GSEA of D28 vs D5 found only cell-cell adhesion as an enriched process, while multiple processes related to early neuronal specification, such as cell projection and axonogenesis, were negatively enriched **[Additional File 1: Fig S3a and Additional File 3: Table S2c]**. GSEA of host genes of circRNAs decreased at D28 vs NES did not identify any enriched pathways. However, among Cluster 2 circRNA hosts, terms associated with morphogenesis and development were overrepresented **[Additional File 1: Fig S3b and Additional File 3: Table S2d]**. Additionally, we examined the enrichment of neurodevelopmental disorder (NDD) risk genes since circRNA expression has previously been found to be disrupted in individuals with ASD and SCZ. The tested genes included ASD genes from the Simons Foundation Autism Research Initiative (SFARI) database, collated SCZ risk genes **[Additional File 4: Table S3]**, and developmental disorder-related genes covering broader phenotypes (Li *et al*, 2022a). We found significant enrichment of ASD genes among hosts genes of circRNAs increased at D28 vs NES, and significant enrichment of SCZ risk genes in Cluster 4 circRNA hosts **[Additional File 1: Fig S3c,d]**.

**Figure 2.**
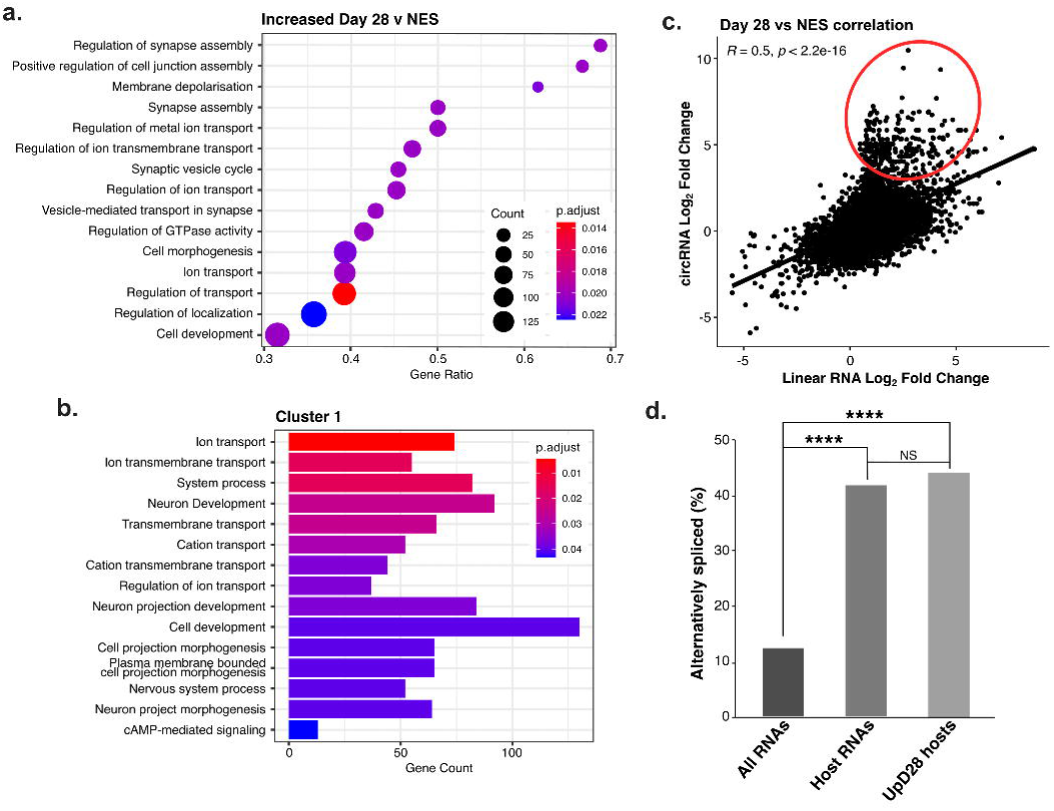
CircRNAs increased during NES differentiation arise from host genes with synaptic function. **a.** GSEA of gene ontology – biological process terms among the genes that host circRNAs increased at D28 of differentiation compared to NES. **b.** Most highly enriched gene ontology – biological process terms among host genes of Cluster 1 circRNAs. **c.** Correlation of LFC values for circRNAs and their corresponding linear counterparts. **d.** Alternative splicing comparison between all RNAs, RNA which host circRNAs and RNAs that host circRNAs increased at D28 vs. NES. **** = p <0.0001, hypergeometric p-value test.

Employing the same RNAseq dataset, we performed differential expression analysis of linear RNAs to identify regulatory patterns and gene ontology terms. The expression of linear RNAs was found to follow a similar regulatory pattern to that of circRNAs, with a higher proportion of linear RNAs increased at D28 vs NES (2,904 genes compared to 1,363 decreased; p-adjusted <0.05, LFC ≥|2|) **[Additional File 1: Fig S4a,b and Additional File 5: Table S4a]**. GSEA also revealed enrichment of similar pathways related to synaptic function among linear RNAs increased by differentiation **[Additional File 1: Fig S4c and Additional File 5: Table S4b]**. Since multiple studies have found discordances between the changes in circRNA levels and the changes in the levels of linear RNAs from the same host gene, we compared changes in circRNA and linear counterparts. As shown, a moderate but significant positive correlation was detected (R = 0.5, p <2.2e-16). Plotting of LFC correlates highlighted this same positive linear association; however, there was an evident shift from the linear distribution suggesting that highly increased circRNAs had higher fold-changes than their linear counterparts [**Fig 2c]**. A direct comparison of circRNAs to linear RNA counterparts found this same trend among the top 20 circRNAs with an average increase of circRNA expression of ∼6.7 fold higher than linear RNAs **[Additional File 1: Fig S5]**. Additionally, among circRNAs increased at D28 vs NES, a large majority (71.6%) were over two-fold higher than their linear counterparts from the same host genes, supporting a non-linear relationship between the expression of these two RNA forms.

To further explore the relationship between linear RNA and circRNA regulation, we analysed differential exon usage in our dataset using DEXseq. All exon bins detected by DEXseq were analysed as exons, including those corresponding to circRNA-forming exons. Comparing cells differentiated up to D28 with NES cells we identified a large number of differentially used exons (19,502 exons; p-adjusted <0.05, lfc ≥|2|). Among these, the number of exons with elevated expression was over three times higher than exons with reduced expression (4,680 down:14,822 up), implying substantially increased exon inclusion at D28. This corresponds with alternative splicing of 4,904 RNAs with an average of ∼31 exons. When enrichment of gene ontology terms among alternatively spliced RNAs were examined, pathways related to the ribosome and spliceosome were most significantly enriched **[Additional File 1: Fig S4d]**.

To establish if alternative splicing is associated with circRNA biogenesis and/or increased abundance of circRNAs at D28, the rate of alternate splicing (based on differential exon usage at D28 vs. NES) was compared among three groups: all linear RNAs, linear RNAs sharing host RNAs with circRNAs, and linear RNAs sharing host RNAs with circRNAs increased at D28. While alternative splicing was ∼3-fold higher among RNAs which host circRNAs (p < 0.0001) compared to all RNAs, there was no difference in alternate splicing rate between all host RNAs and hosts of circRNAs increased at D28 (p = 0.153) **[Fig 2d]**. Similarly, we found no correlation between circRNA fold-changes and estimated exon fold-changes (R = -0.018), suggesting that increased abundance of circRNAs during NES differentiation is independent of general splicing of host precursor RNAs.

### Variant frequency is increased in circRNA-forming exons

To further assess the role of circRNAs in human health and disorders, we examined genetic variant burden in circRNA-forming exons. Exons were classified into circ and nonCirc groups based on all exons of the longest annotated transcript containing the back-spliced exon(s). CircRNA exons were further differentiated as exons that either form BSJs or do not form BSJs (nonBSJ exons) **[Fig 3a]**. Genetic variants included in the Genome Aggregation Database (gnomAD) were initially utilised to examine the frequency of circRNA exon variation in the general population. Rates of genetic variation were found to be slightly higher in both circ (p = 2.2e-16) and BSJ (p = 1.4e-5) exons when compared with nonCirc and nonBSJ exons, respectively **[Fig 3b]**. However, no shift was observed in the annotation of variant impact among exon groups **[Fig 3c]** or in the proportion of missense variants or the proportion of missense variants identified as deleterious **[Additional File 1: Fig S6a,b]**. When we limited the analysis to splice variants, a higher rate among circRNA-forming exons in comparison to nonCirc exons was detected (p < 2.2e-16). Furthermore, among circRNA-forming exons, the rate of splice variants was significantly lower in BSJ exons compared with nonBSJ exons (p = 1.1e-4) **[Fig 3d]**. These observations suggest circRNA-forming and BSJ exons are subject to increased variation while splicing variants are less frequent among back-spliced exons.

**Figure 3.**
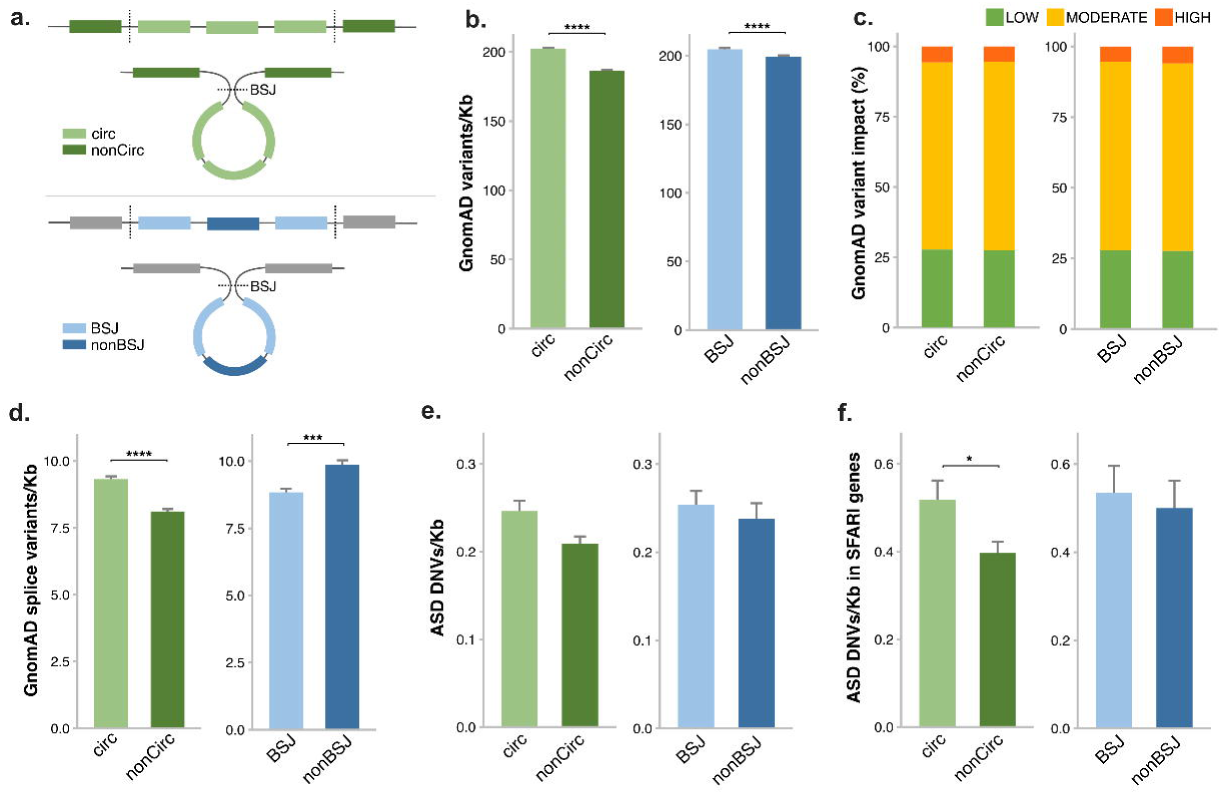
Genetic variant frequency in circRNA-forming exons**. A.** Schematic depicting how exons were subdivided for comparative analysis. BSJ = back-spliced junctions. **B.** Frequency of GnomAD exon variants for circRNAs detected in this study, normalised for exon size (Kb). **C.** Proportion of GnomAD exon variants annotated as having either low, medium, or high impact. **D.** Frequency of GnomAD splice-region exon variants/Kb. **E.** Frequency of rare *de novo* ASD variants/Kb. **F.** Frequency of rare *de novo* ASD variants/Kb in exons of circRNAs arising from ASD-associated SFARI host genes. Two sample Z-test/Wilcox test, **** = p < 0.0001, *** = p < 0.001 * = p < 0.05. DNVs = *de novo* variants.

Next, we examined the frequency of rare *de novo* variants found in ASD from a large exome sequencing study from 21,219 family-based samples, including 12,166 variants in 6,430 individuals with ASD (Satterstrom *et al*, 2020). We identified similar results as for gnomAD variants in that *de novo* ASD variants occurred at a slightly higher rate in circ and BSJ exons **[Fig 3e]**. This increase in circ exon variants was significant specifically when examining circRNAs formed from ASD-associated SFARI host genes (p = 0.0384) **[Fig 3f]**. We note however that as genetic variation overall was increased in circularised exons, that this may not be indicative of specific enrichment of ASD-associated variants in circRNAs. Similarly, a trend for increased rare *de novo* variants in BSJ above non-BSJ exons was also detected (p = 0.06017) among 2,179 unaffected sibling controls from the same cohort **[Additional File 1: Fig S6c,d]**.

Due to a lack of a comparable large resource for SCZ, we collated SCZ variants reported from several sources **[Additional File 1: Table S5]**. In total, 2,035 rare *de novo* variants identified in individuals diagnosed with either Schizophrenia or schizoaffective disorder were included. When comparing the frequency of SCZ variants, we found no differences across exon groups either among all transcripts or when comparing variants specifically in SCZ risk genes **[Additional File 1: Fig S6e,f]**.

### Binding site enrichment in circRNAs differentially expressed during NES differentiation

Given that circRNA functionality remains largely unknown, differentially expressed circRNAs were screened *en masse* for potential function by searching for binding sites among circRNA sequences. The discovery that the highly abundant circRNA, *CDR1as,* functions in sequestering miR-7 through numerous seed sites (Memczak *et al*., 2013; Piwecka *et al*, 2017) has since led to frequent reporting of predicted circRNA:miRNA:mRNA interactions (Chen *et al*., 2020; Dell’Orco *et al*, 2021; Liu *et al*, 2021; Ma *et al*, 2019). Therefore, we first investigated miRNA sponging by exploring possible interactions between miRNAs and DEcircs. We limited our search to 125 human miRNAs previously associated with neurodevelopment or NDDs, which were most of interest to our study **[Additional File 6: Table S6a,b]**. To construct putative circRNA:miRNA:mRNA interaction axes, binding sites for these miRNAs were predicted in both DEcircs as well as 3’UTRs of differentially expressed RNAs **[Additional File 6: Table S6c,d]**. Based on canonical miRNA regulation where miRNAs repress gene expression and circRNA sponges alleviate target repression, circRNAs with predicted miRNA bindingsites were matched to mRNAs containing binding sites for the same miRNA and with the same pattern of regulation. We found no circRNAs to have a high number of binding sites for any particular miRNA with a maximum of three sites identified in any single circRNA. This is consistent with other reports that enrichment of miR-7 sites in CDR1as is a unique phenomenon among circRNAs (Guo *et al*, 2014; You *et al*., 2015). We report identified circRNA:miRNA:mRNA interactions in **Additional File 6: Table S6e,** however, we do not expect that these circRNAs function in miR-sponging.

Next, we examined the enrichment of RNA-binding protein (RBP) motifs among DEcirc sequences. A number of motifs were found to be enriched among different subsets of DEcircs, including two motifs enriched among increased circRNAs **[Additional File 7: Table S7a]**. These two motifs were associated with two splicing factor RBPs, namely, TIA1 Cytotoxic Granule Associated RNA Binding Protein Like 1 (TIAL1) and Splicing Factor Proline and Glutamine Rich (SFPQ) **[Fig 4a,b]**. These results highlighted a possible functional link between TIAL1 and SFPQ RBPs and the increased abundance of circRNAs in neuronal differentiation.

**Figure 4.**
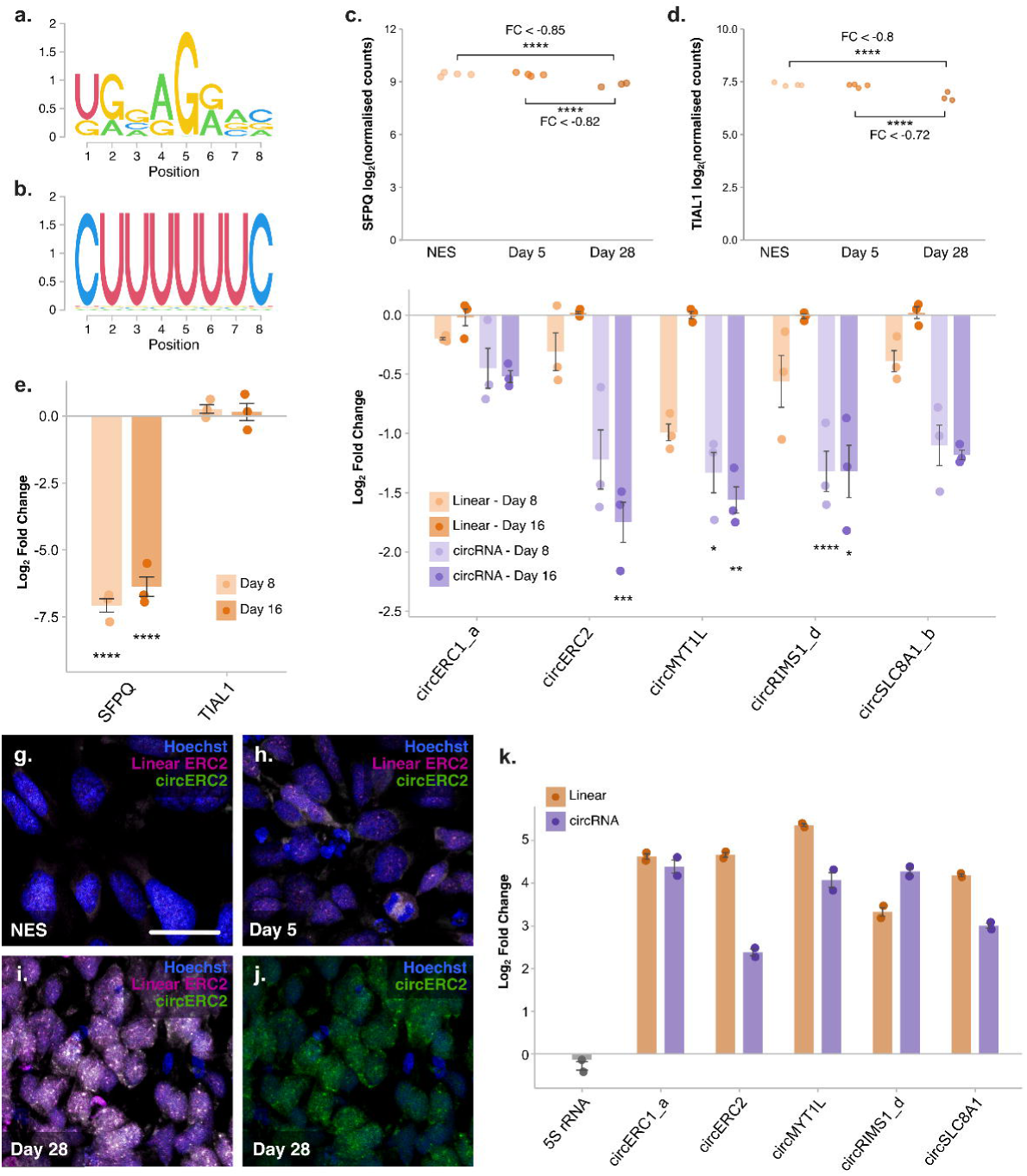
RNA-binding protein SFPQ binds and regulates subsets of circRNAs during neuronal differentiation**. a.** SFPQ motif enriched among circRNAs increased at D28 v D5. **b.** TIAL1 motif enriched among circRNAs increased at D28 v NES. **c.** Normalised counts of SFPQ RNA during NES differentiation. **d.** Normalised counts of TIAL1 RNA during NES differentiation. FC = LFC from DEseq2 analysis, **** = DEseq2 adjusted p-value < 0.0001. **e.** Detection of SFPQ and TIAL1 mRNA following siRNA treatment (*n* = 3). **f.** Relative expression of SFPQ targets following SFPQ knockdown (*n* = 3)**. g-j.** *In situ* detection of *circERC2* and its linear isoform in undifferentiated NES cells (**g**), and cells at D5 (**h**) and D28 (**i,j**) of differentiation. Blue = Hoechst nuclear counter stain, magenta = linear ERC2 probe, green = *circERC2* probe. Scale bar in g = 25 μm, applies to g-j. images are representative of staining from biological replicates (*n* = 3). **k.** SFPQ RIP followed by RT-qPCR analysis to identify SFPQ targets and *5S* rRNA control, normalised to IgG RIP samples from D28 differentiated neurons (*n* = 2). **** = p < 0.0001, *** = p < 0.001, ** = p < 0.01, * = p < 0.05, one-way ANOVA, Tukey post-hoc analysis, Bonferroni adjusted. Error bars represent standard error.

### SFPQ regulates expression of circRNAs in NES differentiation

To further test the relationship between increased circRNAs and enriched RBP motifs, expression of SFPQ and TIAL1 mRNA in NES differentiation was first established. Normalised counts from RNAseq analysis indicated that both RBPs were expressed at all time points albeit with SFPQ levels around tenfold higher than TIAL1 **[Fig 4c,d]**. We also noted a significant reduction in expression of both SFPQ and TIAL1 at D28, although the log-fold reduction was low, this pattern is reflective of SFPQ expression in mice, which is highest during early neurodevelopmental stages and later diminishes (Takeuchi *et al*, 2018). From motif analysis of circRNA sequences, a number of ‘top’ putative targets for SFPQ and TIAL1 were subsequently chosen based on motif frequency, base mean expression and LFC values **[Additional File 7: Table S7b,c]**. Elevated expression of all selected circRNA targets during NES differentiation was confirmed using divergent primer qPCR with all targets increased ∼4 fold or greater as early as day eight of differentiation (D8) **[Additional File 1: Fig S7a]**. To determine if these RBPs affected target circRNA expression, we knocked down SFPQ and TIAL1 during NES differentiation with cells treated with siRNA on days zero (NES) and eight (D8) and RNA collected on D8 and D16. For quantification of protein levels, cell lysates were also collected four days after each siRNA treatment **[Additional File 1: Fig S7b]**. Quantification of mRNA at D8 and D16 found *SFPQ* to be robustly reduced by siRNA treatment compared to non-targeting control siRNA (siNTC) **[Fig 4e]**. SFPQ protein levels were also reduced ∼50% at D4, however, protein expression was returned to baseline levels at D12 indicating that only transient knockdown of SFPQ was achieved **[Additional File 1: Fig S7c,d]**. TIAL1 protein expression was too low to be detected in any samples and *TIAL1* mRNA levels also appeared unaffected by siRNA treatment **[Fig 4e and Additional File 1: Fig S7c,d]**. TIAL1 and its predicted targets were therefore excluded from further analyses.

Following siRNA-SFPQ treatment, we quantified the relative expression of select circRNA targets alongside their corresponding linear mRNA form. Knockdown of SFPQ resulted in depletion of all circRNA targets at both D8 and D16 with an apparent greater decrease of circRNAs compared to linear counterparts, an effect that was especially pronounced at D16 of differentiation **[Fig 4f]**. This indicates that SFPQ affects expression of both circRNAs and their linear counterparts, but we expect that changes in circRNA expression are exacerbated by their enhanced stability (Jeck *et al*., 2013; Memczak *et al*., 2013), even beyond D8 when SFPQ protein was no longer suppressed. To further explore this relationship between SFPQ and circRNA expression we examined SFPQ expression by qPCR at additional timepoints of differentiation **[Additional File 1: Fig S8a]**. Consistent with RNAseq data, we found SFPQ expression reduced at later timepoints of differentiation when circRNA expression is highest. Although this appears counterintuitive for a regulator of circRNA formation, when we further profiled selected SFPQ circRNA targets throughout differentiation up to D50 we found circRNA abundance increases linearly during early differentiation (D8) and plateaus during later stages **[Additional File 1: Fig S8b-f].** Next, as a qualitative analysis, two circRNAs that exhibited the greatest decrease following SFPQ knockdown, *circERC2* and *circMYT1L* were selected for *in situ* hybridisation. circRNA probes were designed to hybridise to the back-spliced junction along with a control probe designed to a linear splice junction of one back-spliced exon. No specific signal was seen with a negative scrambledCirc *in situ* probe or for *circMYT1L*, likely due to low expression as linear MYT1L was only detected at low levels **[Additional File 1: Fig S9a,b]**. However, *circERC2* was detected with a clear increase in expression observed following differentiation **[Fig 4g-j]**. *In situ* detection of *circERC2* showed outspread co-localised with linear ERC2 and expression throughout the neuronal soma **[Fig 4g-j]**. This localisation pattern also appeared unchanged by siRNA-SFPQ treatment **[Additional File 1: Fig S9c-f].** Finally, to determine if SFPQ can bind to the predicted circRNA targets, RNA immunoprecipitation followed by qPCR analysis (RIP-qPCR) was performed on neurons at D28 of differentiation. We found that SFPQ immunoprecipitation enriched all circRNA targets greater than four-fold and, in the case of *circRIMS1_d*, specific enrichment of the circRNA exceeded that of the linear isoform **[Fig 4k]**. Collectively, this suggests a role for SFPQ not only in the regulation of circRNAs during neuronal differentiation but also as a potential binding partner of increased circRNAs.

### CircRNAs increased during differentiation are flanked by long introns

While the SFPQ targets examined were selected based on enrichment for a singular SFPQ motif, additional motifs for SFPQ are known and could be involved in the effects of SFPQ knockdown. Additionally, SFPQ has previously been shown to have an important role in circRNA biogenesis through interactions with circRNA flanking introns, specifically regulating circRNAs with long flanking introns that contain distal inverted Alu repeat elements (IAE) (Stagsted *et al*, 2021). The observed decrease of circRNA targets following SFPQ knockdown could be due to a more general effect of SFPQ promoting circRNA formation through flanking intronic regions. We therefore tested flanking introns of increased circRNAs for enrichment of RBP motifs. Although four motifs for SFPQ were found to be enriched in flanking introns, this represented a small minority among a total of 349 enriched motifs from 76 RBPs, the majority of which were motifs corresponding to members of the HNRNP (93) and SRSF (91) protein families **[Additional File 7: Table S7d]**. Previously, SFPQ binding sites have been identified as enriched both in close proximity (+/- 2000 bp) to BSJ sites of circRNAs (Stagsted *et al*., 2021) as well at 5’ ends of long introns from SFPQ-bound genes (Takeuchi *et al*., 2018). The frequency of SFPQ motifs in flanking introns of circRNA subsets was consequently examined to determine if any position-based pattern was evident. We observed that the frequency of SFPQ motifs among flanking introns of DEcircs is highest at intron ends distal to the BSJ and lowest in regions closest to the BSJ **[Fig 5a]**. Analysing motif position relative to the 5’ site of both upstream and downstream flanking introns indicated that this pattern was driven by an enrichment of SFPQ binding sites at 5’ intronic sites **[Fig 5b]**. While this indicates that 5’ SFPQ motifs may be important for biogenesis of circRNAs differentially expressed in this dataset, this pattern was not unique to increased circRNAs and, in fact, circRNAs decreased during differentiation appear to have the highest frequency of 5’ SFPQ motifs in their downstream flanking introns.

**Figure 5.**
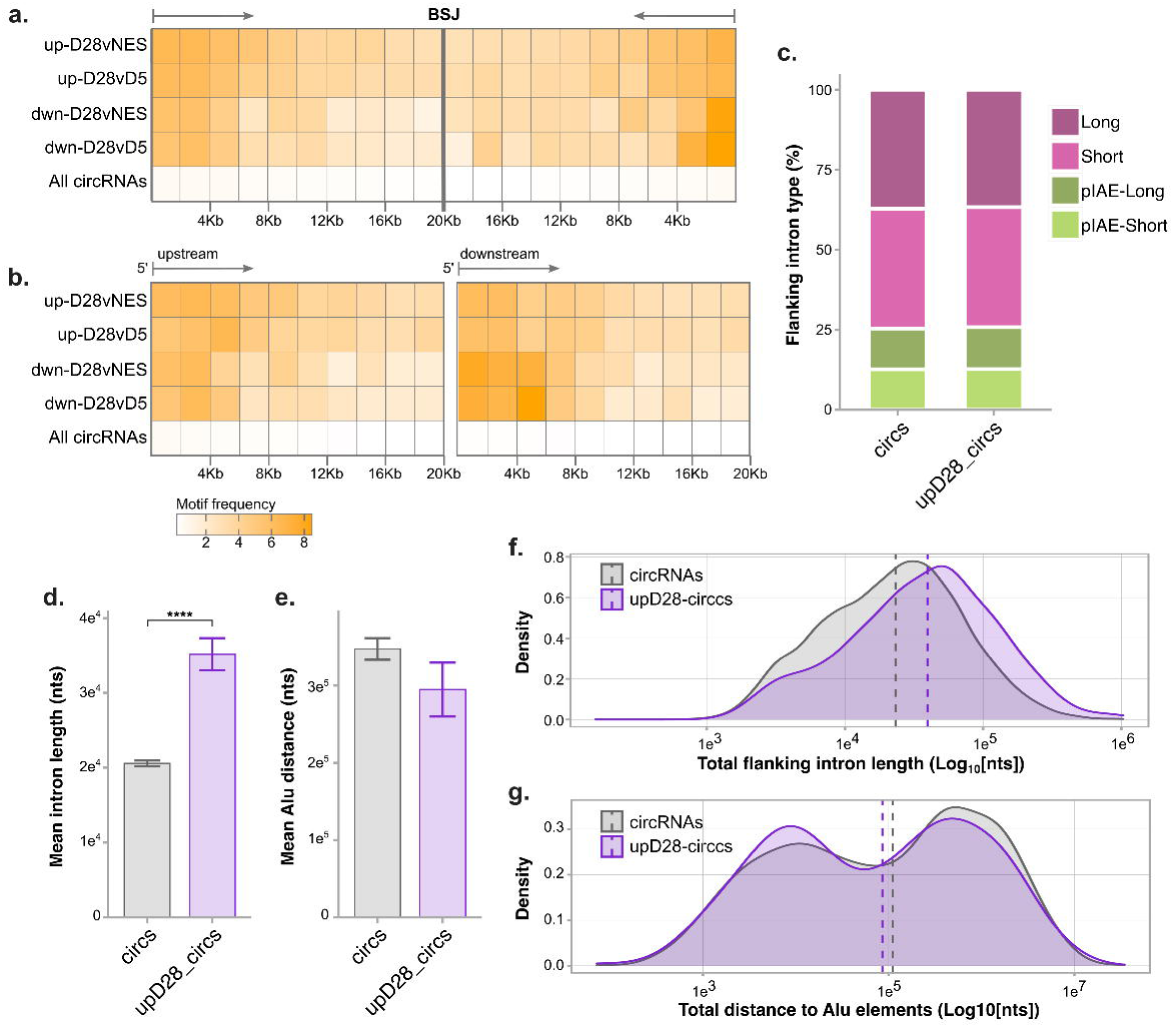
Long introns flank circRNAs increased by NES differentiation. **a.** Frequency of SFPQ motifs in circRNA flanking introns by distance relative to intron splice site distal from the BSJ. **b.** Frequency of SFPQ motifs in circRNA flanking introns by distance relative to the 5’ intron end. Scale in b applies to a and b. **c.** Proportion of flanking intron types among all circRNAs compared to increased circRNAs. Intron types were defined as either, short/long introns with a proximal IAE (pIAE-short/pIAE-long) or short/long introns without proximal IAE (Short/Long). **d.** Mean intron length of flanking introns. **e.** Mean distance to IAE from BSJ. **f.** Density plot of intron length totalled from both flanking introns. **g.** Density plot of IAE distance totalled from Alu element distance on both flanking introns. Error bars in d and e represent standard error, dashed lines in g and f indicate median values. Two sample Z-test, **** = p < 0.0001.

We further compared both flanking intron length and the location of inverted Alu repeats in all circRNAs and circRNAs increased at D28 as potential features of SFPQ-regulated circRNAs. Contrasts were made only between introns flanking circRNAs as, consistent with prior studies (Ivanov *et al*, 2015; Jeck *et al*., 2013), we found flanking introns were significantly longer overall when compared to all introns in circRNA host genes **[Additional File 1: Fig S10a-d]**. Based on grouping of flanking introns by length and presence of proximal IAEs, the proportion of intronic types was unchanged in the subset of increased circRNAs **[Fig 5c]**. A comparison of mean length, however, found that flanking introns were significantly longer in increased circRNAs (p = 2.6e-11) whereas distance to Alu elements in circRNAs that contained IAEs did not differ significantly (p = 0.1615) **[Fig 5d-g]**. When this analysis was repeated on decreased circRNAs we found both mean intron length and Alu to be unchanged **[Additional File 1: Fig S10e-h]**. These results suggest that intronic length, but not IAE distance, may be an important factor in the increased abundance of circRNAs during NES differentiation.

## Discussion

Despite mounting evidence highlighting an important role for circRNA regulation in normal brain function, our understanding of circRNA generation and function during neurodevelopment remains poorly understood. Our study provides a comprehensive profile of circRNA expression in a human model of neuronal differentiation. Although previous reports have suggested high conservation of circRNAs, more recent evidence indicates that circRNA exons exhibit similar conservation to neighbouring linear exons and that the majority of circRNAs are species-specific with only ∼20% of mouse circRNAs having human homologs (Gruhl *et al*, 2021; Guo *et al*., 2014; Santos-Rodriguez *et al*, 2021). Exploring circRNA expression and function within a human-genetic context is therefore critical to understanding the role of circRNAs in both disease and normal brain development.

Recent expansions of the human circRNAome also coincide with the creation of various bioinformatic tools for the detection of circRNAs from sequencing of either total RNA or RNA enriched for circRNA by RNase treatment (Hansen, 2018; Sharma *et al*, 2019). Sequencing of circRNA enriched samples offers improvements in detecting lowly expressed circRNAs and the advantage of reconstructing complete circRNA sequences (Rybak-Wolf *et al*., 2015). By contrast, lowly expressed circRNAs are unlikely to be captured in total RNAseq data, and detection of circRNAs, which relies only on BSJ reads, is prone to false positives. However, the pipeline employed in our study, demonstrates that the identity and expression levels of endogenous circRNAs can be reliably determined from total RNAseq by intersecting results of multiple tools, at least for abundant circRNA species that are most likely to be biologically relevant.

An overall pattern of increased circRNA expression following neuronal differentiation has previously been observed in studies of rodent neuronal development, with elevation of circRNAs occurring following neuronal maturation *in vitro* and within the developing mouse hippocampus (Rybak-Wolf *et al*., 2015; You *et al*., 2015). Enrichment of GO terms related to neuronal and synaptic functions among hosts of increased circRNAs in our dataset is also consistent with several reports highlighting neuronal genes, and especially synaptic factors, as hosts for numerous circRNAs (Ashwal-Fluss *et al*, 2014; You *et al*., 2015). Identification of negatively enriched GO terms related to early, but not late differentiation (cell projection/axonogenesis) at D28 vs D5, however, indicates that circRNAs with altered expression arise from host RNAs functioning in biological pathways relevant to the stage of neuronal maturation. We also observed greater relative increases in the levels of circRNAs than in the levels of linear counterparts, supporting prior evidence of specific circRNA regulation during neuronal development. Non-specific accumulation of circRNAs, however, has been reported in non-proliferating cells and with aging in *C. elegans* (Bachmayr-Heyda *et al*, 2015; Cortes-Lopez *et al*, 2018) while brain-specific circRNAs accumulate during aging in more complex organisms like *Drosophila* and mouse (Gruner *et al*, 2016; Westholm *et al*, 2014). Dissecting the mechanisms of both specific circRNA regulation as well as circRNA accumulation will therefore be crucial for understanding circRNA functions in the developing and aging brain.

Although mechanisms of circRNA biogenesis are still poorly understood, BSJ formation has been shown to occur co-transcriptionally and appears closely linked to linear mRNA splicing (Ashwal-Fluss *et al*., 2014). Differential exon usage has been reported to be in competition with exon circularisation (Ashwal-Fluss *et al*., 2014), whereas other studies identified circRNAs formed primarily from exons with high inclusion rates (Gokool *et al*., 2020). While we observed a significant increase of alternative splicing among transcripts that host circRNAs, alternative splicing was comparable among host RNAs of increased circRNAs, and no correlation was found between circRNA fold-changes and fold-changes of circularised exons. Pathways enriched among differentially spliced RNAs also diverged from those pathways enriched among host precursors giving rise to circRNAs, the latter featuring processes associated with the ribosome and the spliceosome.

Since the majority of individual circRNAs remain unstudied, examining variants occurring in circRNA exons could provide insight into their function and possible roles in evolutionary brain development and disorders. Based on gnomAD data, circRNA exons had slightly higher variant frequency while splice site variant frequency in BSJ exons was significantly lower than nonBSJ exons. This increase in variation was reflected in a higher frequency of rare *de novo* ASD variants in circularised exons of SFARI genes. A separate study also found an enrichment of ASD GWAS SNPs among all circBase circRNAs as well as an enrichment of GWAS SNPs for four out of ten other traits tested (Rajcsanyi *et al*, 2022). Additionally, numerous SNPs which effect circRNA abundance (circQTLs) have been identified in humans with the majority found to be independent, linked only to circRNA not mRNA expression (Ahmed *et al*, 2019). Further, over 600 independent circQTLs have recently been associated with 389 circRNAs specifically in the ASD brain (Mai *et al*, 2022). Taken together, this indicates that circRNA-forming exons in humans may be subject to increased genetic variation and alterations of circRNA function and/or circRNA expression may be an important consideration when determining effects of variants associated with ASD.

While there is mixed evidence of RBP binding site enrichment in circRNAs (Guo *et al*., 2014; You *et al*., 2015), we identified significant enrichment of limited motifs among subsets of DEcircs. SFPQ, found enriched among increased circRNAs, is associated with various aspects of gene regulation and mRNA processing (Heyd & Lynch, 2010; Kaneko *et al*, 2007; Peng *et al*, 2002; Rosonina *et al*, 2005). Though typically localised to nuclear paraspeckles (Passon *et al*, 2012) non-nuclear SFPQ pools, linked to distal axonal transport of RNA granules, have been identified in neuronal cells (Fukuda *et al*, 2021; Furukawa *et al*, 2015; Thomas-Jinu *et al*, 2017). Analogously, SFPQ has been associated with multiple neurodegenerative disorders (Ishigaki *et al*, 2020; Younas *et al*, 2020) and loss of SFPQ causes gross defects in mouse brain development (Takeuchi *et al*., 2018). These effects are associated with defective transcription of long genes, many of which are selectively expressed in the brain, enriched for synaptic functions, and associated with neurodevelopmental disorders (Gabel *et al*, 2015; King *et al*, 2013; Takeuchi *et al*., 2018). A recent analysis in human cell lines also found that SFPQ knockdown primarily depleted circRNAs with long flanking introns containing distal Alu elements, as opposed to short introns with proximal Alu elements (Stagsted *et al*., 2021). Although complementary sequences such as Alu repeats can facilitate circRNA production (Jeck *et al*., 2013; Westholm *et al*., 2014), long flanking introns rather than flanking proximal inverted repeats have been shown to be more important in biogenesis of the most abundant and conserved human circRNAs (Stagsted *et al*, 2019; Westholm *et al*., 2014).

We observed that SFPQ appears to be an important player in regulation of circRNAs and, consistent with previously examined SFPQ targets (Stagsted *et al*., 2021), introns flanking circRNAs increased by NES differentiation were significantly longer than introns flanking all detected circRNAs, while Alu element distance did not appear to be a defining characteristic. Depletion of circRNAs following SFPQ knockdown during differentiation indicates that SFPQ is involved in circRNA biogenesis in our model, possibly through previously described mechanisms such as 5’ end enriched SFPQ binding in long flanking introns. However, motif enrichment in increased circRNA flanking introns indicates many other RBPs were enriched to a greater extent, and patterns of intronic SFPQ motif enrichment appeared relatively uniform across all DEcircs. This finding indicates that, while SFPQ is likely a critical factor in circRNA biogenesis, additional factors other than intronic SFPQ binding must be involved in directing specific circRNA regulation. Given that SFPQ was uniquely enriched within increased circRNA sequences, we speculate that these exonic binding sites may also play a role in circRNA regulation. Previously, in addition to intronic sites, exonic sites were shown to contribute to circRNA biogenesis from *musclebind like splicing regulator 1* (*MBLN1*) pre-mRNA through binding of its own protein product, MBLN1, thereby regulating its own expression (Ashwal-Fluss *et al*., 2014). Here, we show that multiple circRNA targets were also enriched by SFPQ immunoprecipitation, suggesting SFPQ binding sites in these circRNAs are occupied and could function in sequestering SFPQ. Further research is needed to untangle the functional significance of SFPQ binding in circRNAs elevated during differentiation either in regulating their own biogenesis, altering host gene splicing and/or sequestering of SFPQ protein.

## Conclusions

As increasing evidence points to a significant role for circRNAs in neurodevelopment, our data provides an important resource for future studies. We evaluate alternative splicing and differential abundance of linear RNAs and circRNAs and scrutinise genetic variation as well as RBP binding sites, providing an in-depth characterisation of circRNAs in human neuronal differentiation. We also discerned further correlates between circRNAs and neurodevelopmental disorders and identified the splicing factor SFPQ as an important player in neuronal maturation, as both a regulator of circRNA levels and possibly a target itself of circRNA regulation. We note, however, that these mechanisms are not mutually exclusive and a more complex regulatory picture, encompassing splicing modulation, circRNA biogenesis and SFPQ:circRNA binding, is likely yet to be resolved.

## Methods

### NES cell culture and differentiation

Established neuroepithelial stem (NES) cells, previously derived from iPSCs established from a male neurotypical donor, CTRL-9-II; RRID:CVCL_JL74 (Uhlin *et al*, 2017), were seeded on tissue culture flasks coated with 20 μg/ml poly-L-ornithine (Sigma-Aldrich P3655), and 1 μg/ml laminin (Sigma-Aldrich L2020). Cells were grown in DMEM/F12+GlutaMAX medium (ThermoFisher 31331093) supplemented with 0.05X B27 (ThermoFisher 17504044), 1X N2 (ThermoFisher 17502001), 10 ng/ml bFGF (fisher scientific CTP0261), 10 ng/ml EGF (PeproTech AF-100-15) and 10 U/ml penicillin/streptomycin (ThermoFisher 15140122). Medium was exchanged 50% daily and cells maintained in 5% CO_2_ at 37°C, passaging once 100% confluent and seeding at a density of 5×10^4^ cells/cm^2^. Neural differentiation was induced by growth factor withdrawal the day after plating with media B27 concentration increased to 0.5X. Media was exchanged 50% every second day up until D15, after which media was supplemented with 0.4 ug/ml laminin and exchanged 50% every three days.

### RNA extraction and RNA sequencing

Cells were lysed in TRIzol reagent (ThermoFischer 15596026) before separating with chloroform and mixing the aqueous phase with isopropanol as per manufacturer directions. RNA was then isolated from the isopropanol/chloroform solution using the ReliaPrep RNA Cell Miniprep kit (Promega Z6010). For linear RNA depletion, RNA was incubated with 20 U RNase R (Lucigen RNR07250) for 30 mins at 37 °C, as described previously (Pandey *et al*, 2019), before proceeding with the same RNA extraction protocol. For RNAseq, three to five biological replicates were extracted for each cell line and time point. Libraries were prepared with Illumina Truseq Stranded total RNA RiboZero GOLD kit and sequenced on the NovaSeq6000 platform with a 2×151 setup using NovaSeqXp workflow in S4 mode flowcell. Raw reads were processed using cutadapt v3.2 to trim adaptor sequences and low-quality base pairs and discard short reads (options: -m 20 -e 0.1 -q 20 -O 1). The GRCh37 genome assembly was used for all alignment, annotation, and downstream analysis steps.

### CircRNA detection from RNAseq

Back-spliced junctions were detected from paired end reads using three separate programs; CIRCexplorer2 v2.3.8 (Zhang *et al*, 2016), MapSplice2 v2.1.8 (Wang *et al*, 2010) and CIRI2 v2.0.6 (Gao *et al*, 2018). For CIRCexplorer2, modules were run with default options, reads were pre-aligned with TopHat v2.0.9 tophat_fusion (with Bowtie v1.1.2 and Samtools v0.1.19) using –fusion-min-dist 200. MapSplice was run using python v2.7.6 and options –fusion-non-canonical –min-fusion-dist 200. CIRI2 was run using Perl v5.26.2 with default settings after aligning reads with BWA v0.7.17 (with Samtools v1.9). The R package circRNAprofiler v1.4.2 (Aufiero *et al*, 2020) was used as described in the manual to merge and filter back-spliced junction reads from all three detection programs as well as for circRNA annotation and extraction of circRNA sequences.

### Differential expression and gene ontology analyses

Based on clustering analysis, two outlier samples (1xNES/1xD5) were initially removed before differential expression analysis of both circRNAs and genes using DESeq2 v1.30.1 and applying the ashr method of LFC shrinkage (Stephens, 2017). Thresholds of LFC values ≥ |2| and p-adjusted <0.05 were used for sub-setting circRNAs and genes as differentially expressed. For analysis of expression over all three time points, results from DESeq2 likelihood ratio test were analysed with DEGreport v1.26.0. Analysis of differential exon usage was performed using the DEXSeq package (v1.40.0) (Anders *et al*, 2012) using provided python scripts to generate a flattened annotation file of exon bins, with aggregation disabled, and to count reads within exon bins. The standard analysis pipeline described in the tutorial was followed, defining exon bins as differentially used if estimated LFC was ≥|2|, padj <0.05 and exon base mean >10 and classifying genes as differentially spliced if at least one exon of a gene was differentially used. As some exon bins defined by DEXSeq differed from annotated exon regions, differential exon usage data was compared with circRNA exons based on position information and using the intersect function of bedtools software v2.29.2. For annotated exons which contained multiple exon bins, exon fold-changes were calculated as the mean estimated exon bin fold-change. The clusterProfiler package v3.18.1 was used for all gene ontology analyses using either biological process terms annotated in org.Hs.eg.db v3.12.0 or user input annotations for analysis of NDD-associated gene enrichment. As GSEA requires only one entry per gene and numerous host genes had multiple circRNAs, data were split into negatively and positively regulated circRNAs and the minimum and maximum circRNA LFC values were assigned to host genes with duplicate entries.

### PCR, quantitative PCR and Sanger sequencing validation of circRNAs

For all experiments, cDNA was generated using the iScript cDNA synthesis kit (Bio-Rad). To validate circRNA detection, divergent primers **[Additional File 1: Fig S1b]** were designed as previously described (Panda & Gorospe, 2018). PCR was performed using HotStarTaq (Qiagen) and bands visualized by gel electrophoresis. For Sanger sequencing, sequences were amplified using HotStarTaq and circular primers **[Additional File 1: Fig S1b]**. Products were isolated using DNA Clean and Concentrator-5 kit (Zymo Research) or, if multiple products, bands were separated using gel electrophoresis and extracted using Zymoclean Gel DNA Recovery Kit (Zymo Research). Samples were sequenced by Eurofins Scientific before comparing alignments to predicted annotations with nucleotide BLAST and visualizing alignments with ApE v2.0.49.0. Divergent primers and SYBR green (Bio-Rad) were used for quantitative PCR (qPCR) analyses. For differential expression validation, ciRNA expression was normalised to two reference circRNAs, *circRTN4* and *circCASK1G3,* found to have high and consistent expression throughout NES differentiation. For SFPQ targets circRNAs were normalized to *U6* snRNA. For siRNA experiments all genes/linear RNAs were normalised to *GAPDH* mRNA. SPAG9 and CPSF6 were used as reference genes for profiling SFPQ expression across differentiation time-points. For known circRNAs, divergent BSJ primers were designed using circInteractome (Dudekula *et al*, 2016). For novel circRNA divergent BSJ junctions and for all circRNA primers, circRNA sequences were recreated in SnapGene Viewer v5.1.1 based on annotations from circRNAprofiler and exon sequences in Ensembl GRCh37 and primers designed using NCBI primer blast. All primer sequences are listed in **Additional File 1: Table S8**.

### Variant analysis

For circRNAs which annotated to known splice sites, exon regions were extracted based on the longest transcript containing the upstream and downstream back-spliced exons and sub-grouped as in **Fig 3a**, with the assumption that no exons were spliced out of circRNAs. GnomAD GRCh37 exomes files v2.1.1 were downloaded for each chromosome and converted into bed format. *De novo* variants associated with ASD cases and controls were sourced from Satterstrom *et al*. (2020) and SCZ variants were pooled from various sources listed in **Additional File 1: Table S5**. All variants were intersected with exon subgroups using bedtools and all downstream analysis was performed in R, normalising all exon variant frequencies by exon length. SIFT annotations for gnomAD exon variants were predicted using Ensembl VEP v104.

### miRNA and RBP binding site prediction

A list of 125 miRNAs associated with neurodevelopment or NDDs **[Additional File 6: Table S6a,b]** was collated from literature review and pubmed searches for miRNA/microRNA with terms such as neurodevelopment/neuronal/neuron differentiation etc. Ensembl biomart was used to extract 3’UTR sequences for DEGs and, if multiple 3’UTR sequences were annotated to a single gene, the longest 3’UTR sequence was used. miRNA binding sites in circRNA and 3’UTR sequences were predicted using remote RNA22 v2 with the following parameters: seedmismatchesallowed = 0; seedsize = 8; minpairmatchs = 12; minenergy = -12; maxguwobbles = 1; threshold = 7000000. To examine enrichment of RNA-binding motifs in circRNAs, separate fasta files of DEcircs were generated for circRNAs decreased or increased at D28vNES / D28vD5 and D5vNES as well as all 4 LRT clusters. RNA-binding protein motifs were downloaded from the ATtRACT database v0.99β. Motif enrichment was tested using the AME function of MEME-suite v5.3.0 with default settings and a background dataset of all annotated (5839/6385) circRNAs. For individual motif site detection, the MEME-suite FIMO function was used with a threshold p-value of 0.001.

### siRNA mediated silencing and protein detection

For siRNA mediated knockdown, Accell Human SMARTpool siRNAs targeting SFPQ (E-006455-00-0010) or TIAL1 (E-011405-00-0010) were added to cell media at a final concentration of 0.3 μM. Accell non-targeting control pool (D-001910-10-20), transfected at the same concentration was used as a control treatment (siNTC). Protein samples were collected 96 hr after siRNA treatment. After removing media and washing once with PBS cells collected with a cell scraper in a lysis buffer of 50 nM Tris-HCl, 100 mM 100 mM NaCl, 5 mM EDTA, 1 mM EGTA and 1X HALT protease inhibitor cocktail (PIC; Fischer Scientific) and samples sonicated at 36% amplitude for 6X 1 sec. The Qubit Protein Assay Kit (Thermo Scientific) was used to measure protein concentration. Protein was detected using the Simple-Western-JESS capillary western blot system (Bio Techne) multiplexed for total protein and chemiluminescence detection of mouse αSFPQ (1:2000; Sigma-Aldrich WH0006421M2, RRID:AB_1843565) or mouse αTIAL1 (1:10; Santa Cruz sc-398372). The Compass for S.W. software v5.0.1 was used to analyse data and calculate peak area normalised to total protein area.

### In situ hybridisation

Locked nucleic acid (LNA) probes for *in situ* were designed to be 45-55 bps in length with a melting temperature of ∼70-75°C. Probes were designed to be complementary to the BSJ for circRNA detection while linear control probes were designed to target a linear splice junction, within the same transcript, containing one of the BSJ exons. CircRNA probes plus a negative control scrambled circRNA probe labelled with 5’FAM and linear probes labelled with 5’Cy3 were ordered from Integrated DNA Technologies (IDT) as HPLC purified, Affinity Plus DNA oligos and resuspended in TE buffer. Details of all probe sequences are listed in **Table S9**. Cells were plated on glass coverslips coated with 100 μg/ml poly-L-ornithine and 2 μg/ml laminin. *In situ* hybridisation was performed as described previously with some modifications (Zirkel & Papantonis, 2018). Cells were washed with PBS before incubation in fresh fixative (4% PFA, 0.9% NaCl and 5% Glacial acetic acid in UltraPure H_2_O) for 17 min at room temp. Coverslips were washed for 5 min with PBS before adding ice-cold 70% ethanol and storing at least overnight or up to a few weeks at -20°C. After removing from -20°C cells were recovered for 5 min at room temp in PBS, permeabilised for 10 min in 0.5% Triton X-100/0.5% saponin (Sigma-Aldrich S7900) and washed in PBS. Cells were then treated with 1% formaldehyde in PBS for 10 min and washed once in PBS for 10 min. Cells were dehydrated with ethanol at 70, 90 and 100% for 3 min each and air-dried. For each coverslip 15 ng of labelled probe was diluted in 50 μL of hybridisation buffer consisting of 10% Dextran sulfate (Sigma-Aldrich 265152), 0.1% Sodium dodecyl sulfate (Sigma-Aldrich 71736), 30% Ethylene carbonate (Sigma-Aldrich E26258), 20 μg/mL Yeast tRNA (Thermo Fischer AM7119) and 2X SSC. Probes were denatured at 83°C for 5 min and hybridised overnight at 37°C in a box humidified with 2X SSC. Coverslips were washed 3 times in 2X SSC for 10 min at 42°C, followed by two 3-min PBS washes at room temp. Cells were blocked for 30 min in 0.1% Triton X/5% Donkey serum/PBS and counterstained with 1:1000 Hoechst. After two 3-min washes in PBS, coverslips were air-dried for 3-5 min and mounted onto glass slides using Prolong Diamond Antifade Mountant (Thermo Fischer). Images were acquired using an LSM 700 Zeiss Confocal Microscope and contrast was enhanced uniformly.

### Formaldehyde RNA immunoprecipitation

Formaldehyde RNA immunoprecipitation (fRIP) was performed as previously described (Hendrickson *et al*, 2016) with some modifications. NES cells were differentiated until D28 in a T75 flask. To collect, cells were washed with PBS, rinsed with 6 mL Accutase for 10 sec and incubated in 4 mL TrypLE select for 2 min/37°C before addition of 200 μL of Neuronal isolation enzyme (Thermo Scientific, 88285) and further incubating 1 min/37°C. Dissociation was stopped with 6 mL of Trypsin inhibitor and solution collected, any remaining cells were collected by rinsing flasks with 6 mL of Wash Medium (DMEM/F12+GlutaMAX, 5% FBS, 10 U/ml penicillin/streptomycin). Clumps were broken up by gentle pipetting and cells spun at 400 x g for 3 min and resuspended in Wash Medium at 5×10^6^ cells/mL. Crosslinking was performed by addition of formaldehyde to a final concentration of 0.1% and incubation for 10 min with rotation. Glycine was added at a concentration of 125 mM, incubating for 5 min with rotation to stop the reaction. Cells were pelleted, washed twice in cold PBS and once in PIC, and stored at -80°C. For immunoprecipitation, cell pellets were resuspended and lysed by incubation for 10 min at 4 °C with rotation in RIPA lysis buffer (50 mM TRIS [pH 8], 150 mM KCL, 0.1% SDS, 1% Triton-X, 5 mM EDTA, 0.5% sodium deoxycholate, 0.5 mM DTT, 1X PIC, 100 U/mL RNaseOUT). The lysate was sonicated 15 times (0.7s on, 1.3s off, Vibra-Cell VCX-600, Sonics) at an amplitude of 10% and centrifuged at max speed for 10 min at 4°C. Supernatant was collected and diluted with an equal volume of Binding buffer (150 mM KCl, 25 mM Tris [pH 7.5], 5 mM EDTA, 0.5 % IGEPAL) with 0.5 mM DTT, 1X PIC and 100 U/mL RNaseOUT added. Cells were incubated 2 hr at 4°C with rotation after addition of either 6 μg mouse αSFPQ (Sigma-Aldrich WH0006421M2, RRID:AB_1843565)) or 6 μg normal mouse αIgG (Santa Cruz sc-2025, RRID:AB_737182). To each immunoprecipitation, 50 μL of Dynabeads Protein G (Thermo Fischer 10004D), pre-washed 2X in Binding buffer, was added before incubating 1 hr/4°C with rotation. Beads were washed 2X 10 min/4°C with rotation in 1 mL Binding buffer/PIC/RNaseOUT. To reverse cross-linking, beads were resuspended in 56 μL UltraPure water and 33 μL reverse crosslinking buffer (PBS (without Mg or Ca), 6% N-lauroyl sarcosine, 30 mM EDTA, 15 mM DTT), 2 mg/mL Proteinase K and 400 U/mL RNaseOUT were added to each reaction, before incubating 1 hr/42°C followed by 1 hr/55°C. Bead/buffer mix was added to 1 mL TRIzol and RNA extraction performed as described above. For each fRIP, 5 ng of RNA was used for cDNA synthesis and targets were analysed by qPCR, normalizing SFPQ-fRIP to IgG-fRIP.

### Intron analysis

Sequences for introns flanking circRNA back-spliced exons were extracted using BSgenome v1.58.0 and Biostrings v2.58.0 based on annotations output by circRNAprofiler. RBP binding sites were predicted using MEME-suite as described above and SFPQ motif frequency for each subset of circRNAs normalised by the number of circRNAs. All GRCh37 repeat elements were downloaded from UCSC table browser RepeatMasker track and Alu elements extracted using grep. Bedtools intersect was used to subset Alu elements that overlapped with intronic regions by at least 50% of the repeat element length and bedtools closest tool used to find Alu elements closest to the BSJ site. CircRNAs were classified as having inverted Alu elements when the closest Alu elements in the downstream and upstream flanking introns were in opposite orientation (on opposite strands). CircRNAs were classified as having proximal IAE flanking introns based on a total Alu distance from the BSJ less than the median for all circRNAs with IAEs. Classification of circRNAs with short or long flanking introns was also classified based on median length, totalled from both flanking introns.

### Statistics and data visualisation

All statistics and data visualisation were performed in R v4.0.4. Specific statistical tests performed are described within the relevant results section. All replicates specified within figure legends indicate biological replicates i.e., cells cultured separately at successive passages. Various R packages were used for visualisation of data including ggplot2 v3.3.3, EnhancedVolcano v1.8.0 and VennDiagram v1.6.20

## Declarations

### Ethics approval and consent to participate

Not Applicable

### Consent for publication

Not Applicable

### Availability of data and materials

Generated raw sequencing data is unavailable as it contains potentially identifiable human data. All other related data and code are available at: https://github.com/Tammimies-Lab/circRNA_neuro.git

### Ethics approval and consent to participate

Not Applicable

## Authors’ contributions

MW, KT & MG designed experiments. MW, MO, SL & FM performed experiments. MW analysed data and wrote the manuscript. All authors read and approved the final manuscript.

## Acknowledgments

The authors acknowledge support from the National Genomics Infrastructure in Stockholm funded by Science for Life Laboratory, the Knut and Alice Wallenberg Foundation and the Swedish Research Council, the SNIC/Uppsala Multidisciplinary Center for Advanced Computational Science for assistance with massively parallel sequencing and access to the UPPMAX computational infrastructure, and the iPS Core facility at Karolinska Institutet for assistance with the generation of iPSC and NES cells.

## Additional Files

### Additional File 1

Format: Portable document format (.pdf)

Title: Supplementary information

Description: Figure S1. Validation of circRNA expression. Figure S2. Validation of circRNA exon retention. Figure S3. Pathway enrichment among differentially expressed circRNAs. Figure S4. Pathway enrichment among differentially expressed linear RNAs. Figure S5. Log2 fold-change values of top increased circRNAs and their corresponding linear host RNAs (D28 vs NES). Figure S6. Variant analysis in circRNA exons. Table S5. Source studies for compilation de novo variants identified in individuals with Schizophrenia or Schizoaffective disorder. Figure S7. SFPQ and TIAL1 target detection and siRNA knockdown. Figure S8. Extended expression profile of SFPQ and identified circRNA targets in NES differentiation. Figure S9. In Situ detection of circRNAs during NES differentiation. Figure S10. Analysis of introns from circRNA host genes. Table S8. Primer sequences.

### Additional File 2

Format: Excel workbook (.xlsx)

Title: Table S1

Description: Supplementary Table 1. Differential expression analysis of circRNAs in NES differentiation. a. Hotspot host genes which generated over 10 circRNAs. b. Pairwise differential expression (DE) analysis of circRNAs at D28 of differentiation compared to NES. c. Pairwise DE analysis of circRNAs at D5 of differentiation compared to NES. d. Pairwise DE analysis of circRNAs at D28 compared to D5 of differentiation. e. Likelihood ratio test DE circRNA analysis of all time points. f. Clustering of LRT results.

### Additional File 3

Format: Excel workbook (.xlsx)

Title: Table S2

Description: Supplementary Table 2. Gene ontology analysis of circRNAs differentially expressed during differentiation. a. Biological processes enriched among host genes of Cluster 1 cicrRNAs. b. Positive gene set enrichment analysis of D28 compared to NES circRNA host genes. c. Positive gene set enrichment analysis of D5 compared to NES circRNA host genes. d. Biological processes enriched among host genes of Cluster2 circRNAs.

### Additional File 4

Format: Excel workbook (.xlsx)

Title: Table S3

Description: Supplementary Table 3. Collated list of Schizophrenia risk genes. a. Gene list. b. Source studies.

### Additional File 5

Format: Excel workbook (.xlsx)

Title: Table S4

Description: Supplementary Table 4. Analysis of linear RNA expression in NES differentiation. a. Differential expression analysis of linear RNAs at D28 compared to NES. b. Positive gene set enrichment analysis of linear RNAs at D28 compared to NES.

### Additional File 6

Format: Excel workbook (.xlsx)

Title: Table S6

Description: Supplementary Table 6. Prediction of circRNA:miRNA:mRNA interactions. a. Collated list of miRNAs linked to neurodevelopment. b. Reference studies for miRNA list. c. miRNA sites predicted in DEcircs. d. miRNA sites predicted in 3’UTRs of differentially expressed mRNAs. e. circRNA:miRNA:mRNA interactions with correlated expression during differentiation.

### Additional File 7

Format: Excel workbook (.xlsx) Title: Table S7

Description: Supplementary Table 7. Prediction of RNA binding protein (RBP) motifs in circRNAs and thier flanking introns. a. Significantly enriched RBP motifs within sequences of differentially expressed circRNA subsets. b. SFPQ target selection from positive circRNA sequences increased at Day 28 compared to Day 5. c. TIAL1 target selection from positive Cluster 1 circRNA sequences and circRNA sequences increased at Day 28 compared to Day 5. d. Significantly enriched RBP motifs within flanking intron sequences of circRNA increased at Day 28 compared to NES.

