## Additional File 1 for "Circular RNAs arising from synaptic host genes are modulated by SFPQ RNA-binding protein and increased during human neuronal differentiation"

### SUPPLEMENTARY INFORMATION

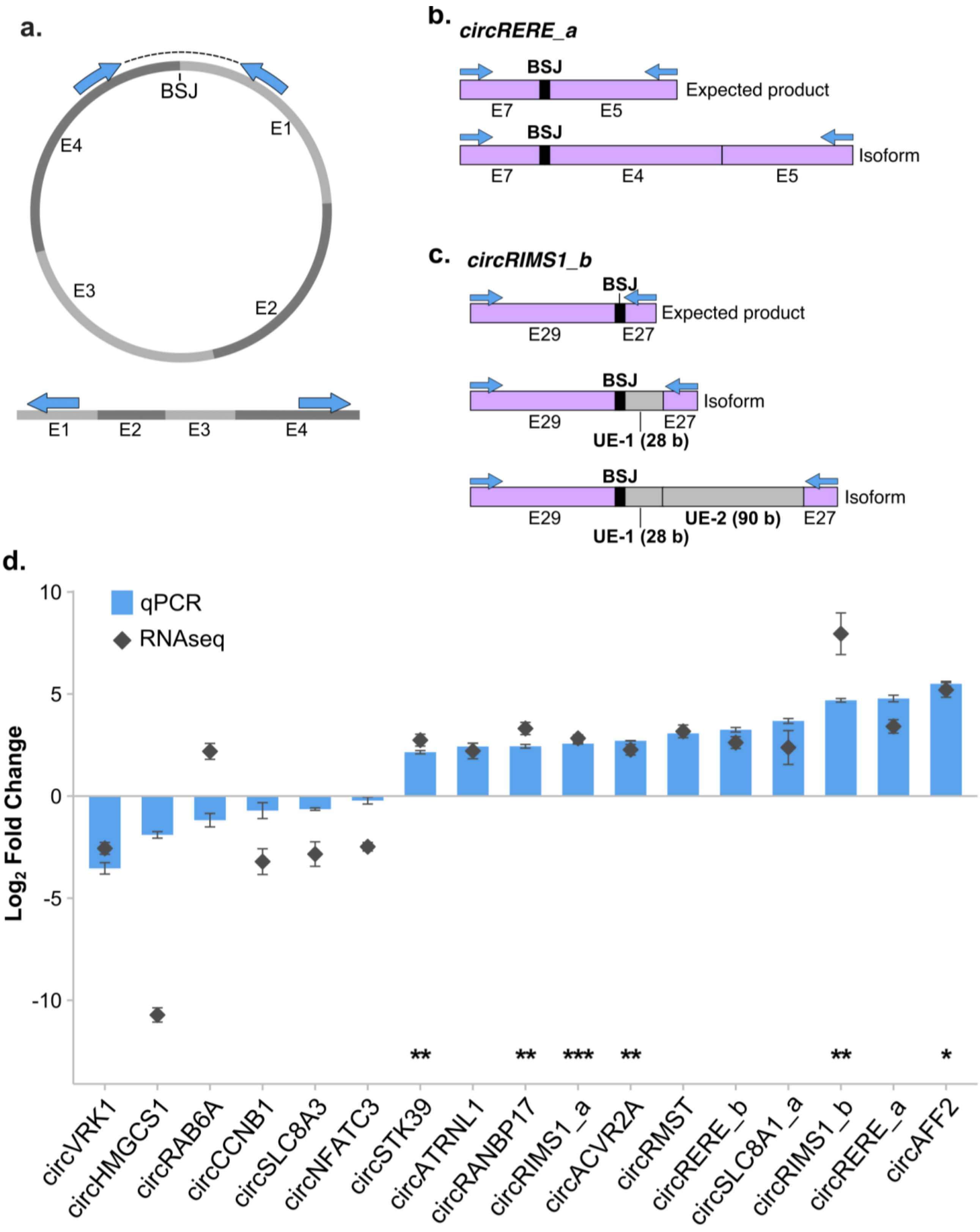

**Figure S1.** Validation of circRNA expression**. a.** Schematic representation of junction primer design (blue arrows) in relation to the circular and linear forms of RNA. **b,c.** Schematic representation of Sanger alignment from isoforms amplified by junction primers for *circRERE_a* (b) and *circRIMS1_b* (c). BSJ = Back Spliced Junction, E = Exon. UE = Unannotated Exon. **d.** Log fold-change of differentially expressed circRNAs from qPCR and RNAseq data, D28 vs NES, *n* = 3, two-sample t-tests, Bonferroni corrected. Error bars represent SEM, ** = p-adj <0.05, ** = p-adj <0.01, *** = p-adj <0.001.

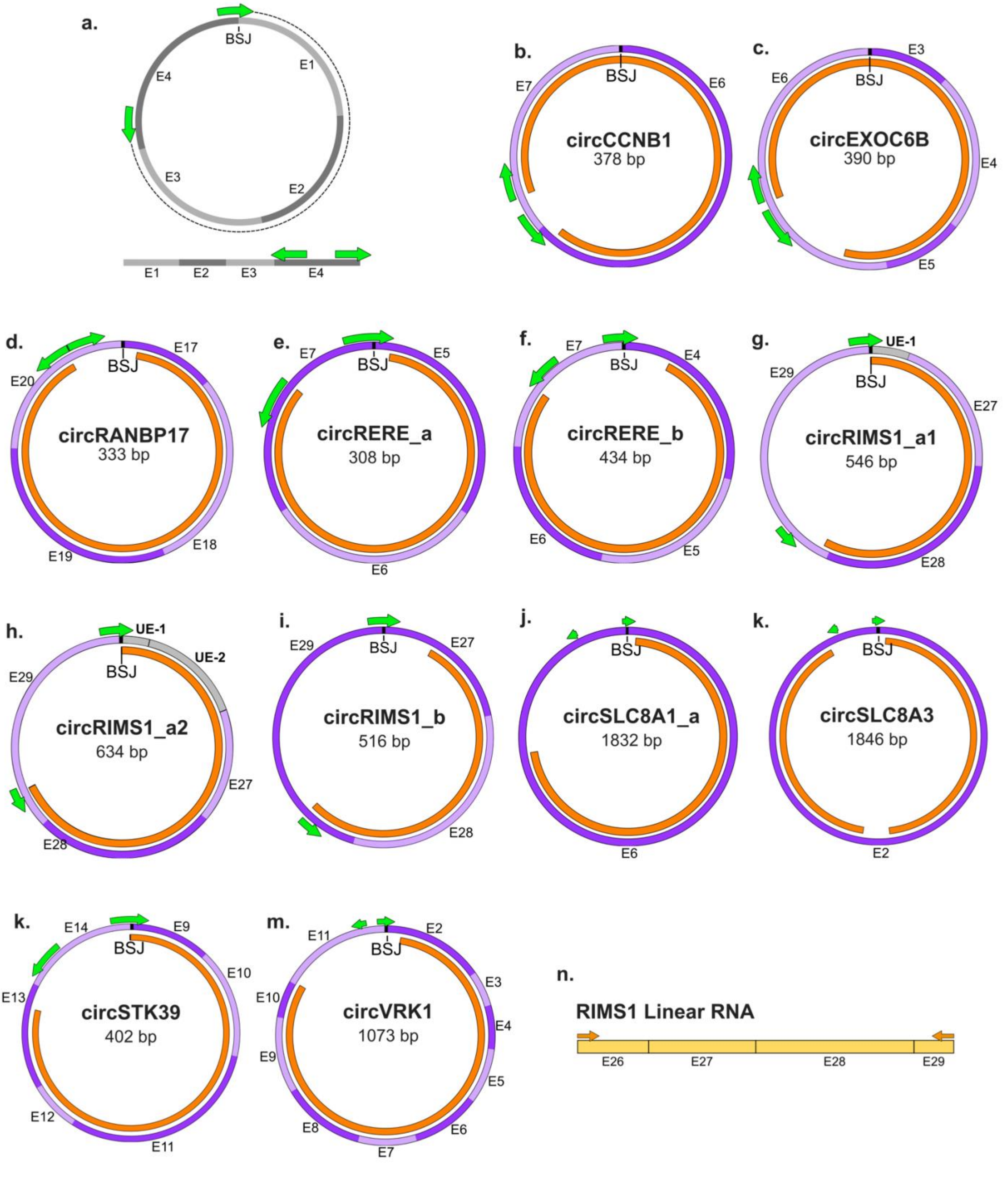


**Figure S2.** Validation of circRNA exon retention. **a.** Schematic representation of circular primer design (green arrows) in relation to the circular and linear forms of RNA. **b-m.** Schematic representations of predicted circRNA exon annotations and sanger sequencing alignments. Exons are shown in purple, green arrows indicate primer location and sanger sequencing product alignment is shown in orange. BSJ = Back spliced Junction, E = Exon, UE = Unannotated Exon. **n.** Schematic representation of Sanger sequencing results from PCR targeting exons 26 – 27 of the linear RNA form of RIMS1, flanking the unannotated exons detected in circular RIMS1 isoforms.

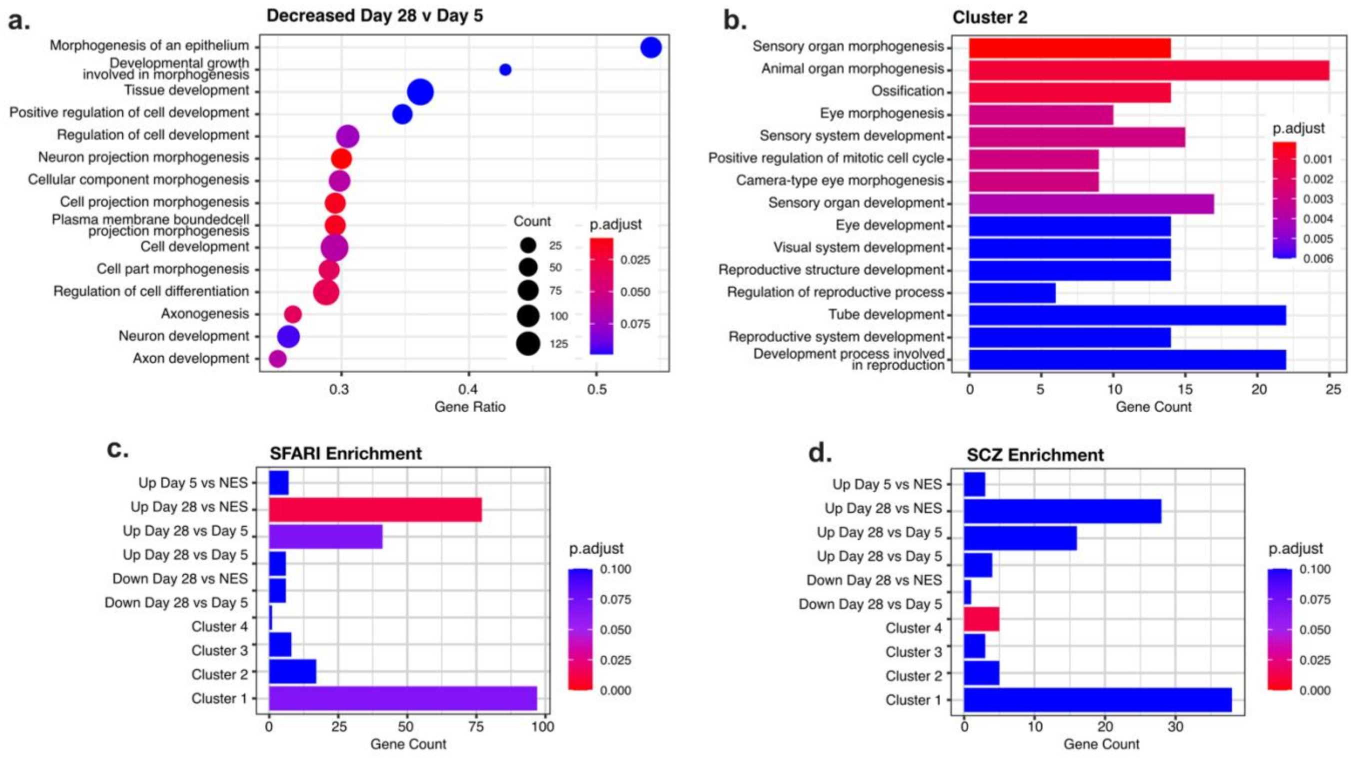


**Figure S3.** Pathway enrichment among host genes of differentially expressed circRNAs. **a.** GSEA of gene ontology - biological process terms among host genes of circRNAs decreased at Day 28 vs Day 5. **b.** Most highly enriched gene ontology - biological process terms among Cluster 2 circRNA host genes. **c.** Enrichment of SFARI risk genes among DEcirc host gene sets. **d.** Enrichment of Schizophrenia (SCZ) risk genes among DEcirc host gene sets.

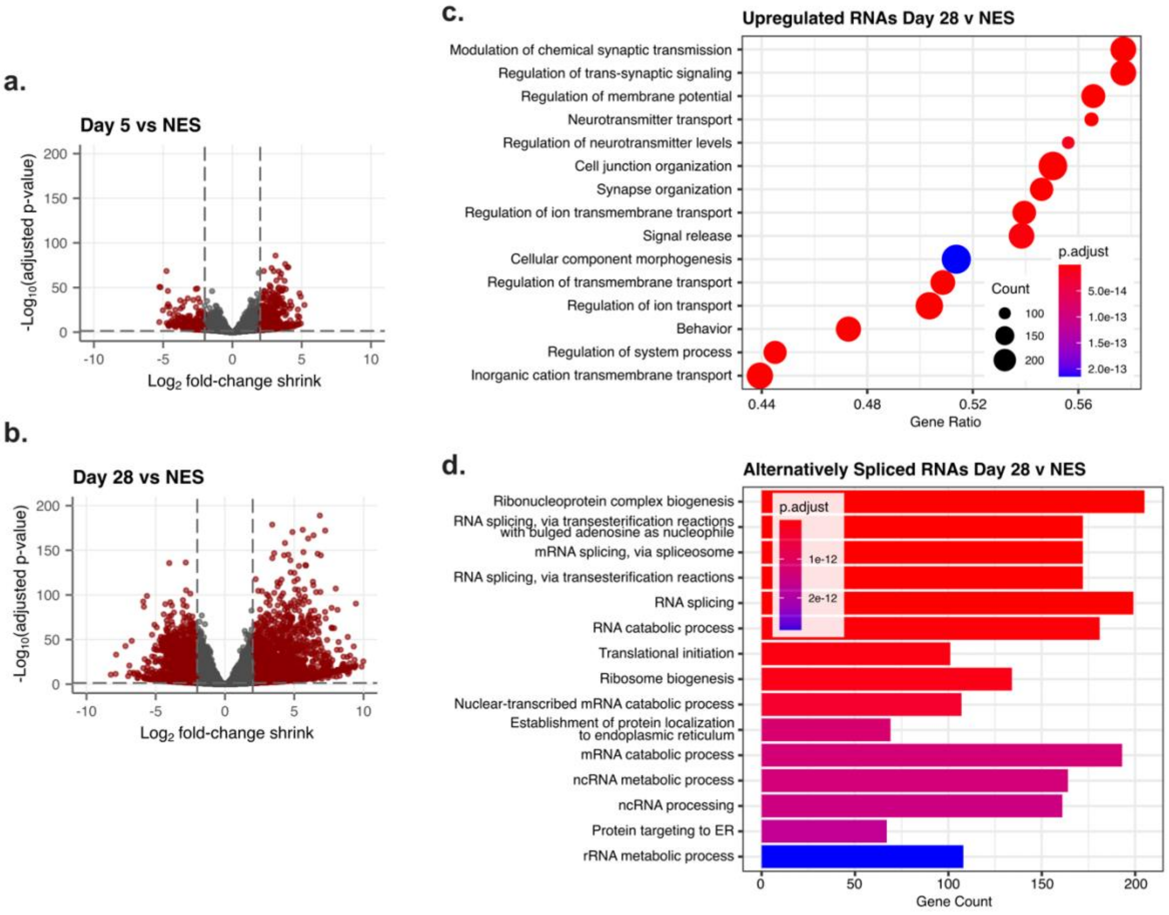


**Figure S4.** Pathway enrichment among differentially expressed linear RNAs. **a,b**. Volcano plots of differentially expressed RNAs at D5 and D28 of differentiation compared to NES. **c.** GSEA of gene ontology - biological process terms among RNAs increased at D28 vs NES. **d.** Most highly enriched gene ontology - biological process terms among RNAs that were alternatively spliced at D28 vs NES.

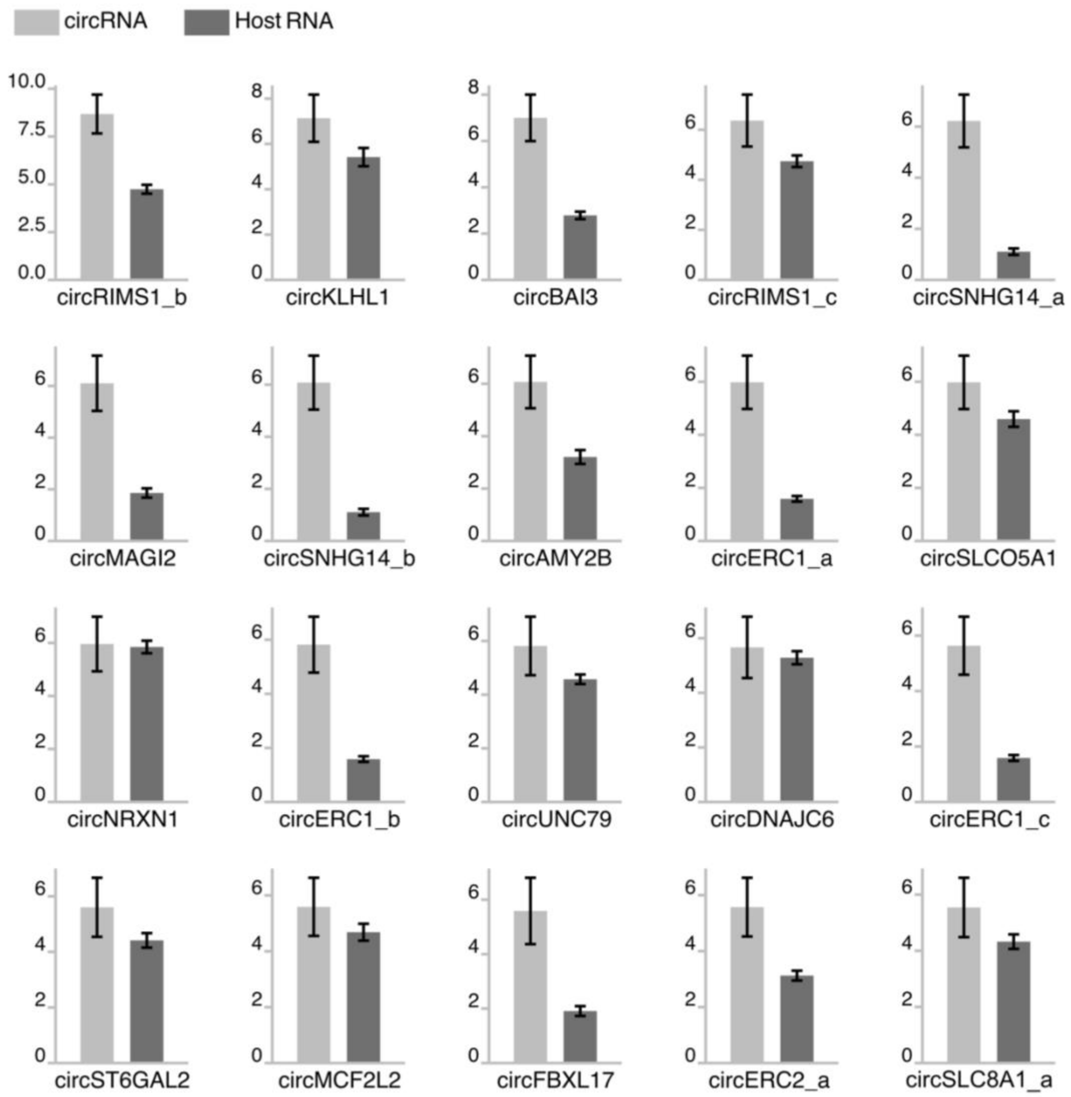


**Figure S5.** Log2 fold-change values of top increased circRNAs and their corresponding linear host RNAs (D28 vs NES).

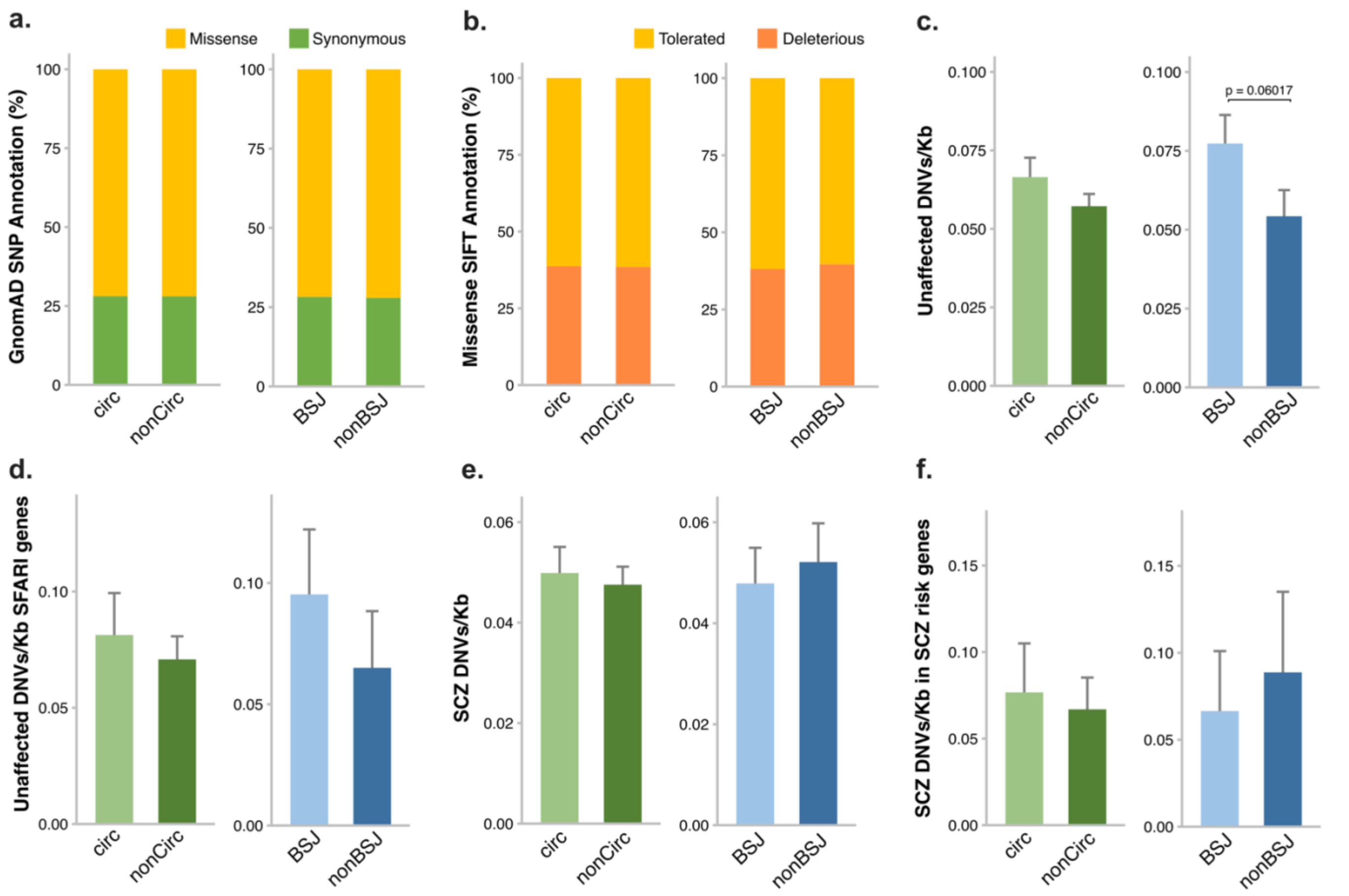


**Figure S6.** Variant analysis in circRNA exons**. a.** Proportion of missense or synonymous SNPs among gnomAD variants. **b.** Proportion of gnomAD missense SNPs classified as 'Tolerated' or 'Deleterious' by SIFT prediction. **c.** Frequency of rare *de novo* variants/Kb in unaffected sibling controls. **d.** Frequency of rare *de novo* variants/Kb in unaffected sibling controls in exons of circRNAs arising from ASD-associated SFARI host genes. **e.** Rate of Schizophrenia (SCZ) *de novo* variants per kilobase in exons of circRNAs. **f.** Rate of SCZ *de novo* variants per kilobase in exons of circRNAs arising from Schizophrenia risk host genes. DNVs = *de novo* variants.

**Table S5.** Source studies for compilation of *de novo* variants identified in individuals with Schizophrenia or Schizoaffective disorder.

| | Authors (Year) | Journal | DOI | Data Type | Gene association with Schizophrenia |
|---|---|---|---|---|---|
| **[1]** | Ambalavanan *et al* (2015) | *Eur. J. Hum. Genet.* | DOI: 10.1038/ejhg.2015.218 | WES | Damaging or missense *de novo* mutations in cases. |
| **[2]** | Fromer *et al* (2014) | *Nature* | DOI: 10.1038/nature12929 | WES | *De novo* mutations in schizophrenia probands (no missense predictions). |
| **[3]** | Girard *et al* (2011) | *Nat. Genetics* | DOI: 10.1038/ng.886 | WES | Damaging or missense *de novo* mutations in cases. |
| **[4]** | Guipponi *et al* (2014) | *PLoS ONE* | DOI: 10.1371/journal.pone.0112745 | WES | Damaging or missense *de novo* mutations in cases. |
| **[5]** | Gulsuner *et al* (2013) | *Cell* | DOI: 10.1016/j.cell.2013.06.049 | WES | Damaging or missense *de novo* mutations in cases. |
| **[6]** | McCarthy *et al* (2014) | *Mol. Psychiatry* | DOI: 10.1038/mp.2014.29 | WES | Damaging or missense *de novo* mutations in cases. |
| **[7]** | Takata *et al* (2014) | *Neuron* | DOI: 10.1016/j.neuron.2014.04.043 | WES | Loss-of-function *de novo* indel variants. |
| **[8]** | Wang *et al* (2015) | *Scientific Reports* | DOI: 10.1038/srep18209 | WES | Nonsynonymous *de novo* variants. |

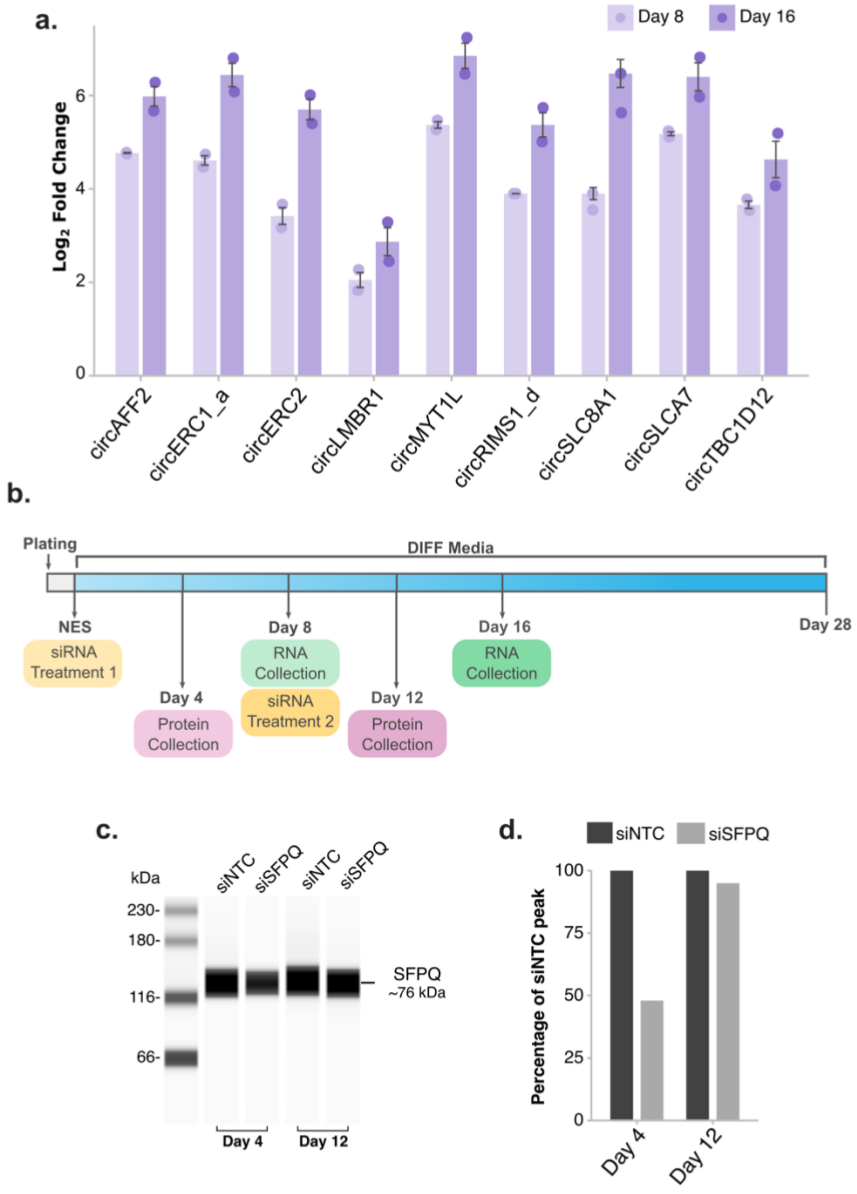


**Figure S7.** SFPQ and TIAL1 target detection and siRNA knockdown**. a.** Detection of circRNA targets during differentiation by qPCR, *n* = 2. **b.** Schematic of siRNA knockdown experiment timeline and sample collection. **c.** Detection of SFPQ protein following siRNA treatment by WES. **d.** Quantification of SFPQ WES, normalised to total protein.

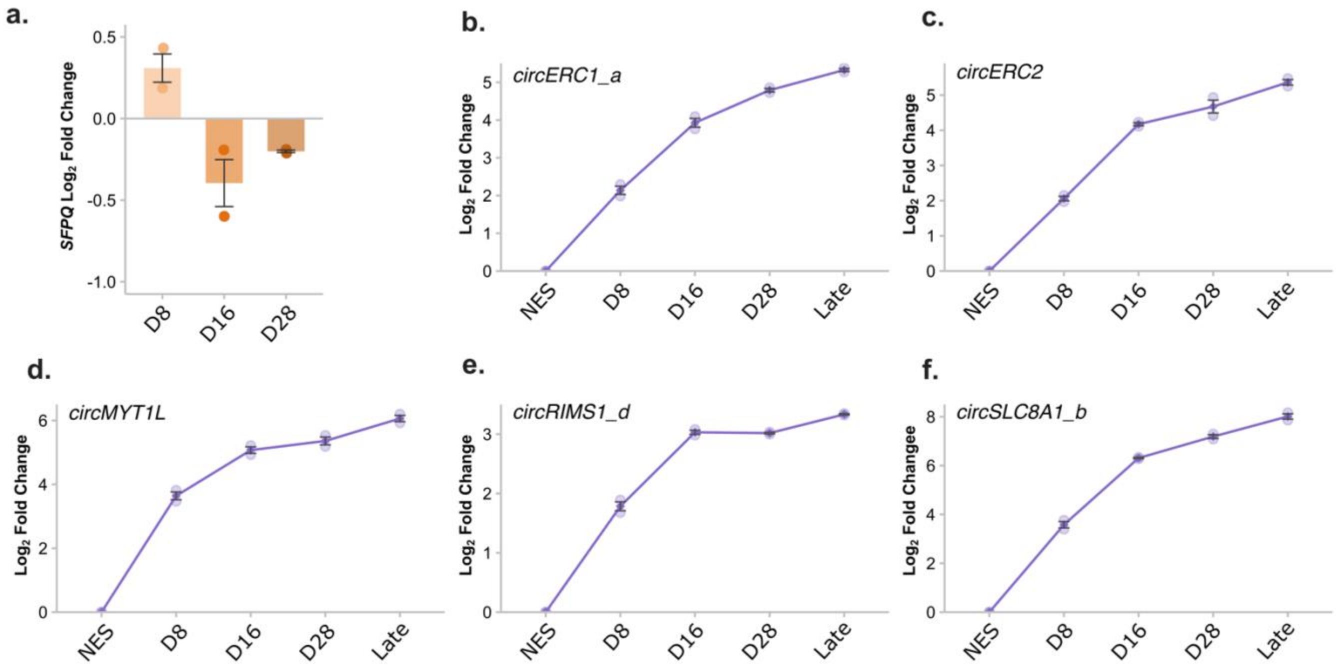


**Figure S8.** Extended expression profile of SFPQ and identified circRNA targets in NES differentiation. **a.** SFPQ expression at days 8, 16 and 28 of differentiation relative to NES cells $n = 2$. **b-f.** SFPQ target circRNA expression detected by qPCR from NES to samples collected after either 40 or 50 days of differentiation (Late) $n = 2$.

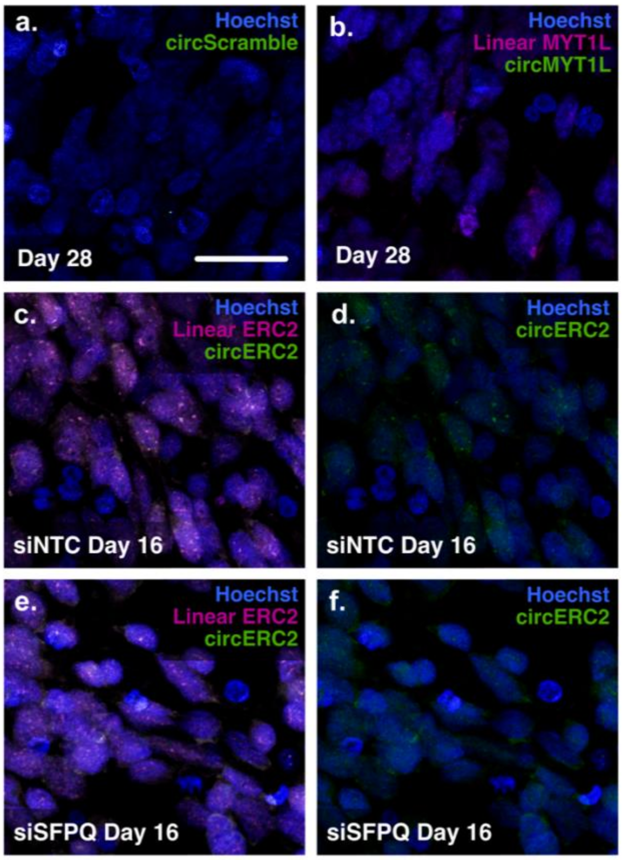


**Figure S9.** *In Situ* detection of circRNAs during NES differentiation. **a.** Signal from the negative control circScramble probe at D28. **b.** Detection of *circMYT1L* and its linear counterpart at D28. **c,d.** Detection of *circERC2* and its linear counterpart at D16 following no-target siRNA control treatment (siNTC). **e,f.** Detection of *circERC2* and its linear isoform at D16 following siRNA knockdown of SFPQ (siSFPQ). Blue = Hoechst nuclear counter stain, magenta = linear probes, green = circ probes. Scale bar in a = 25 μm, applies to all images, images are representative of staining from biological replicates ($n$ = 3).

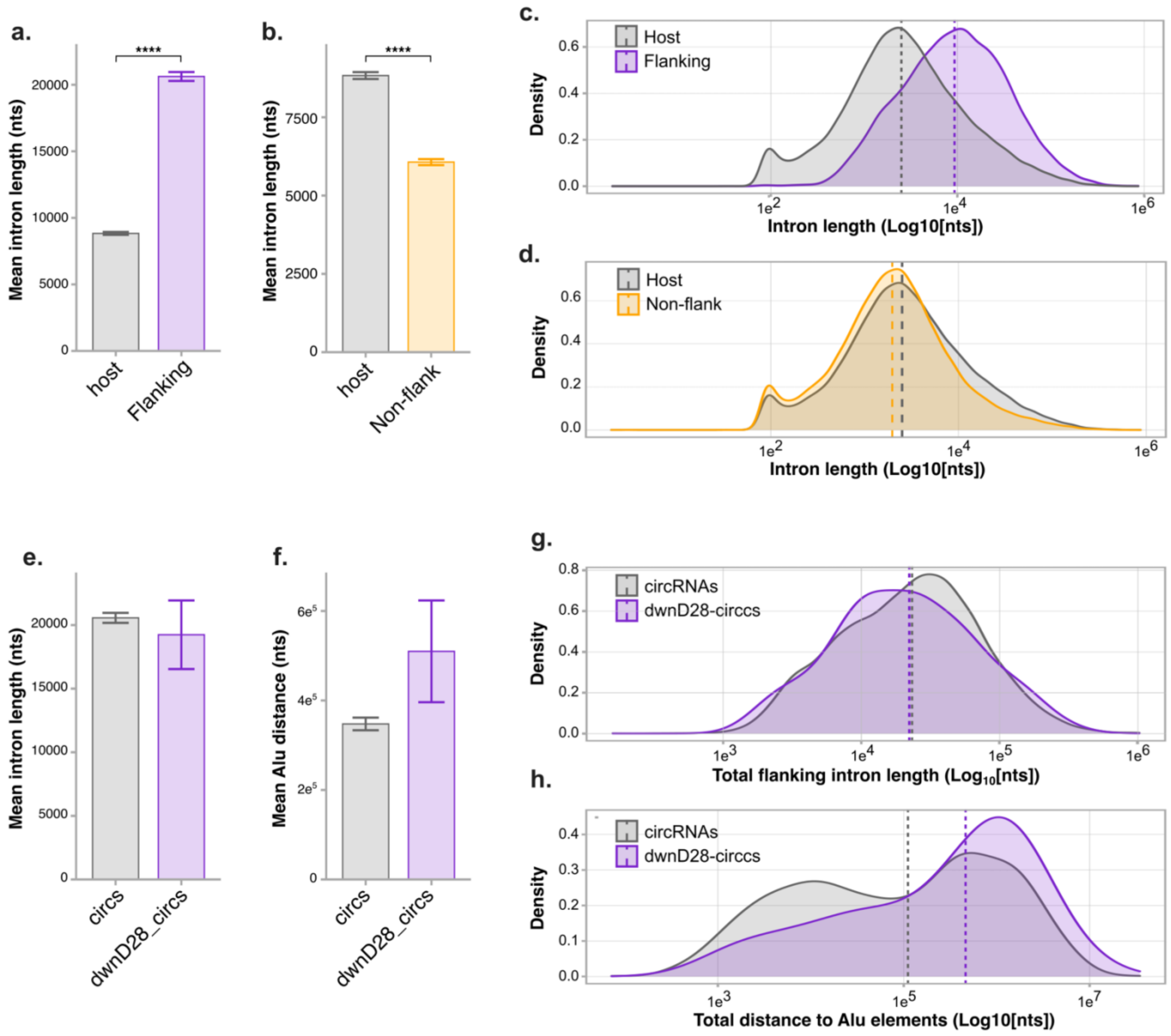


**Figure S10.** Analysis of introns from circRNA host genes. **a.** Mean intron length of circRNA flanking introns compared to all introns in circRNA host genes. **e.** Mean length of host introns that do not flank circRNAs compared to all introns in circRNA host genes. **c.** Density plot of length of circRNA flanking introns compared to all introns in circRNA host genes. **d.** Density plot of length of host introns that do not flank circRNAs compared to all introns in circRNA host genes. **e-f.** Analysis of introns flanking circRNAs decreased at D28vNES compared to all circRNAs. **e.** Mean intron length of flanking introns. **f.** Mean distance to IAE from BSJ. **f.** Density plot of intron length totalled from both flanking introns. **g.** Density plot of IAE distance totalled from Alu element distance on both flanking introns Error bars in a, b, e and f represent standard error, dashed lines in c, d, g and h indicate median values. Two sample Z-test, **** = p < 0.0001.

**Table S8.** Primer sequences.

| Target | Forward primer 5' > 3' | Reverse primer 5' > 3' |
|---|---|---|
| | *Divergent BSJ Primers* | |
| *circACVR2A* | ATTGCGGGGATTGTCATTTG | ACCAGTTTGATTGGTTCTGTC |
| *circAFF2* | GTCATTTGGAACACTCTTGG | GACTCGGTTGGCAAGTG |
| *circATRNL1* | CATTCTATGTGTACGTCAGCAAC | TTTATGGCAGTATGGTGGCGAT |
| *circCCNB1* | GCACTGAAAATTCTGGATAATGGTG | TTGCTCGACATCAACCTCTCCA |
| *circCSNK1G3* | GAATATGTGGCAATTAAGTTGGAGC | ATGTGAGCTGATATTGATAGAGAGC |
| *circERC1_a* | TCTCTTGGAGCTTTCGTCCTC | GGCCATCAGTAACTCCTCCAC |
| *circERC2* | TTGGCCAACCTCCGGATTG | CTGCGCACTTCTTCTAGTAACTG |
| *circEXOC6B* | GGAGAGCATCCGCAAACATTC | TAGTCGACACTGCTTCAGCTC |
| *circHMGCS1* | TCAGCAATTAGCAGGGAAGAG | TGCATATGTGTCCCACGAAG |
| *circLMBR1* | GGTCAGTTGCTAGTGAAGCC | TAAAAGCAAAACAGCCCCAGC |
| *circMYT1L* | AGCCTAAATCCAGTGACAGCC | GCCACCAGTTCATCGTAATTGTC |
| *circNFATC3* | TAGCCGAGGGGCAGTAAAAG | GTGGTAAGCAAAGTGGTGTGG |
| *circPTK2* | CAGCTAGTGACGTATGGATGTT | GACGCATTGTTAAGGCTTCTTG |
| *circRAB6A* | AGGGAGAGAGGAAAGCCAAAGA | GTTCCTGACCCGCAGTATCCC |
| *circRANBP17* | TCGAGGGATTGCCTTTGCAC | TTTTAGCATGAATTTCACAGCATCT |
| *circRERE_a* | AGAGTTTAAAGCCCGAGTGGA | TCGATCCTGAACCAAATGCTGA |
| *circRERE_b* | AGAGTTTAAAGCCCGAGTGGA | GGTCACACAAAGCAGGAGTTG |
| *circRIMS1_a* | AGCAGGTGGAAAGAAACGGA | AGCTTGCTTTTGTGGAAGAGTTA |
| *circRIMS1_b* | AGCAGGTGGAAAGAAACGGA | CGCTCCTCCACTACTAAGC |
| *circRIMS1_d* | TCAGTGATGTTTCCGCCATTTC | TTGTTCGCTCCTCCACTACT |
| *circRMST* | TTGCTCAAGGTGGAACGACA | TTGCTCAAGGTGGAACGACA |
| *circRTN4* | ACGATAAAGGAACTCAGGCG | AATAGGCTGGCACCAAACAC |
| *circSLC4A7* | GATGAAATGGCCAAAACTGCAC | TCCAAGTTTCCAGGAGCAGAC |
| *circSLC8A1_a* | GTGGAGGGGAGGATTTTGAG | TCCCATTGAAAAGGTGGGTGAA |
| *circSLC8A1_b* | TGCACCATTGGCCTGAAAG | GGCGGGGCTCTCCAATCTC |
| *circSLC8A3* | GACAGTAGAAGGGACAGCCA | TTGCAGCACCAGTTGTCCTC |
| *circSTK39* | AGCTTCTTCTTGTGCCGTGA | TCTGGTGTTCTTGTAAGCAGC |
| *circTBC1D12* | GGGGAGGACCTGTAAACCAC | TCCAACCAGGAGCACTATGG |
| *circVRK1* | TGGACCTCAGTGTTGTGGAG | CTCCAACTGCAAATTGTTCTGC |
| *circZNF91* | AGAAGTGTAATTACTGTCAAACGAC | TGCTCTGGCCAAAAGTCTTGA |

| | Circular divergent primers | |
|---|---|---|
| *circATRNL1* | GCTGGAACAATATCTGGGGAAGA | TTGAACCGACAGACCACGTAA |
| *circCCNB1* | GGCCAAATACCTGATGGAACTAACT | TTGCTCGACATCAACCTCTCCA |
| *circEXOC6B* | GGTGGACAACATCCCCAAGC | TTGCAGAATCGATAGTGGCTTAC |
| *circNFATC3* | CAGGGGAAGTGTGACAGAAGATAC | CAGGTGAAGGAACAGGTGAGTG |
| *circPTK2* | GTCTGGATAATCATGGAGCTGTG | TGATGACTCCAATCAGCTTCAC |
| *circRAB6A* | CGTAGCCTCATTCCCAGTTACATCC | GTTCCTGACCCGCAGTATCCC |
| *circRANBP17* | CAAAGACCAGCTACACCATGC | TGTTCAGTGCAAAGGCAATCC |
| *circRERE_a* | ACCCTGAGACAAGAGTAAGAG | TCGGGCTTTAAACTCTCTAGCA |
| *circRERE_b* | CCCTGAGACAAGGTCCACAA | TCGGGCTTTAAACTCTCTAGCA |
| *circRIMS1_a* | CAGCTTAGTCAAACAGAACAAG | GAAGTCCCCATCCGTCTG |
| *linearRIMS1_a* | CAGACACATCGTTCAGCAGTCG | GAAGTCCCCATCCGTCTG |
| *circRIMS1_b* | AGTCAAACAGCAAGCTTAGTAG | GAAGTCCCCATCCGTCTG |
| *circSLC8A1_a* | TGATGAAATTGTTAGGTTGTGACA | CATGATGCCAATGCTCTCAC |
| *circSLC8A3* | ATGATGAAACTGTGTCTCTGGC | CACCTGATGTCCGCAGAAC |
| *circSTK39* | TTGAGATTAAGGCCCACAGCA | TAGTCTTCATTAGCATTGGGTG |
| *circVRK1* | CAATAACAAAGTGAAAATGCCTCGTG | ATTCTCCACAACACTGAGGTC |
| | *Linear Primers* | |
| *5SrRNA* | CGCCCGATCTCGTCTGAT | GGTCTCCCATCCAAGTACTAACCA |
| *AFF2* | CCAGTGGTCAGCCAAACAAG | TGTCTTCCTTTGCAGATGGGT |
| *CPSF6* | GATTCCATGGCATATGGGGC | TGGAGTCTGTCCAGCTTTTAGG |
| *ERC1* | AATCTTCGGGCAGAGAGAAGG | AGAGGACGAAAGCTCCAAGAG |
| *ERC2* | AGCATGGCTGACAACTCACA | TTCTTTTTCGGCCAGGGACT |
| *GAPDH* | AAGGTGAAGGTCGGAGTCAAC | GGGGTCATTGATGGCAACAATA |
| *LMBR1* | CACAGTGATGGGTCAGTTGC | TTAGTCGTCTCTGGAGTGCTTC |
| *MYT1L* | TACGTCATGTTGGGGAAGCC | TTCTGCTGCGGATTCCTCTC |
| *RIMS1* | GGAGGAGCGAACAAGACAGA | GTTCGGGAAATGGCGGAAAC |
| *SFPQ* | CCAGCATGGCACGTTTGAG | TGTCGTCTCATCAGATCTTGGC |
| *SLC4A7* | CTGAACGGATGCTTCAAGATGA | CTTCCACTGTGCAGTTTTGGC |
| *SLC8A1* | CTTGTGGTTGGGACTAACAGC | ACGTAATCGAAACAGGAGGGC |
| *SPAG9* | GCAGCTCATCACCCAGTACG | TCTTCAAGTCTGCTAATCTGGTCA |
| *TBC1D12* | ACATCCAAAATCATTCAGCAGGA | AATCAAGGCAGTGGTAGAAAGC |
| *TIAL1* | TGCGTCTGGGTTAACAGATCA | GGCTGCACTTTCATGGGTTG |

**Table S9.** Probe designs for *in situ* hybridisation. Junction indicates which exon/exon junction the probe spans. Transcript ID = Ensembl transcript ID. Bases highlighted in red indicate bases with locked nucleic acid modifications.

| Probe | Transcript ID | Junction | Sequence 5' – 3' | Length | Tm (°C) |
|---|---|---|---|---|---|
| circERC2 | ENST00000288221 | 14/13 | FAM-TCTTATTCTGATCTTTCATATGCTTCATCTCCAGGATCTCCTCCAG | 46 b | 70.4 |
| Linear ERC2 | ENST00000288221 | 14/15 | Cy3-TGGCTGCAAGTAGTGCTTCCTGTTTCATCTCCAGGATCTCCTCCAG | 46 b | 75.2 |
| circMYT1L | ENST00000428368 | 10/10 | FAM-TGACAATTCATTTGATGGTCTTCATAGTATGGCTTTTTGACATGGCTGTCACTGG | 55 b | 73.1 |
| Linear MYT1L | ENST00000428368 | 10/11 | Cy3-CTCTTTCTTTTCTGTTCTTGAGGGATCATAGTATGGCTTTTTGACATGGC | 50 b | 70.8 |
| Scrambled | - | - | FAM-TGCGTTCTCCTTCTAACTCGTGCAACACTTCGATCTATAGTACTCT | 46 b | 70.3 |
